# Single-cell RNA-sequencing reveals widespread personalized, context-specific gene expression regulation in immune cells

**DOI:** 10.1101/2021.06.04.447088

**Authors:** Roy Oelen, Dylan H. de Vries, Harm Brugge, Gracie Gordon, Martijn Vochteloo, BIOS Consortium, Chun J. Ye, Harm-Jan Westra, Lude Franke, Monique G.P. van der Wijst

## Abstract

Gene expression and its regulation can be context-dependent. To dissect this, using samples from 120 individuals, we single-cell RNA-sequenced 1.3M peripheral blood mononuclear cells exposed to three different pathogens at two time points or left unexposed. This revealed thousands of cell type-specific expression changes (eQTLs) and pathogen-induced expression changes (response QTLs) that are influenced by genetic variation. In monocytes, the strongest responder to pathogen stimulations, genetics also affected co-expression of 71.4% of these eQTL genes. For example, the pathogen recognition receptor *CLEC12A* showed many such co-expression interactions, but only in monocytes after 3h pathogen stimulation. Further analysis linked this to interferon-regulating transcription factors, a finding that we recapitulated in an independent cohort of patients with systemic lupus erythematosus, a condition characterized by increased interferon activity. Altogether, this study highlights the importance of context for gaining a better understanding of the mechanisms of gene regulation in health and disease.

## Introduction

Over a decade of genome-wide association studies (GWAS) has revealed thousands of genetic variants associated with disease risk^1^, most of them single nucleotide polymorphisms (SNPs). Despite this, the cascade of events through which these variants change disease risk remains largely unclear. One way to dissect this cascade is by linking disease-associated SNPs to downstream gene expression through so-called expression quantitative trait locus (eQTL) analysis.^2^ However, recent work by Yao et al. indicated that, on average, only 11% ± 2% of disease heritability is mediated by *cis-*eQTLs, i.e. SNPs affecting the expression of nearby genes.^3^ One explanation for this relatively low contribution could be that many of these eQTL effects are cell type⍰specific and context-dependent^4,5^, which means that their disease contribution cannot be accurately estimated using steady-state expression in bulk-averaged tissues. In other words, the relevant context for a particular disease-associated SNP may not have been studied yet, meaning that many of the true downstream effects of these SNPs remain hidden.^6^ In a first effort to identify tissue-specific eQTLs, the GTEx consortium performed eQTL analysis in 44 different human tissues across 449 individuals (70⍰1361 individuals/tissue).^7^ However, this study was limited by the relatively small number of donors for many of the tissues and the lack of cell type⍰specific resolution. More recently, with the advent of high-throughput, cost-efficient single-cell RNA sequencing (scRNA-seq) technologies^8,9^, it has become possible to assess both the cell type⍰specific and context-dependent effects of risk SNPs on downstream gene expression.^10–12^

While the tissue or cell type is one context that can affect the association between a SNP genotype and gene expression, many other contexts can also be of influence. For the immune system, for example, exposure to specific pathogens commonly occurs and the immune response following exposure can create the environmental context required to change specific interactions between genetics and downstream gene expression.^4,13–17^ In turn, these context-specific interactions may explain why exposure to specific pathogens has been associated with the development of autoimmune diseases in individuals with a genetic predisposition.^18^ For example, a reovirus can disrupt intestinal immune homeostasis and initiate loss of tolerance to gluten in individuals expressing HLA-DQ2 or HLA-DQ8, leading to celiac disease.^19^ Another example is the strong indications that enteroviral infections in the pancreas, such as with coxsackievirus, in genetically predisposed individuals may accelerate the development of type I diabetes (T1D).^20–22^ Several T1D-associated risk genes affect the antiviral response through regulation of type I interferon (IFN) signaling.^23^ When the insulin-producing pancreatic β cells of genetically predisposed individuals are then exposed to such viruses, incomplete viral clearance and chronic infection of these β cells may be the consequence. This could then induce β cell apoptosis that contributes to the development of T1D.^24,25^ Overall, it is estimated that 11⍰30% of autoimmune risk loci involve *cis*-eQTLs in blood, and it is hypothesized that trait-associated eQTLs have increased context-specificity.^26–28^ Given this hypothesized context-specificity, it is important to study eQTLs in a variety of different contexts to determine the possible effect of environment on the interplay between genetic variation and gene expression in disease.

Here we present the 1M-scBloodNL study in which we performed scRNA-seq on 120 individuals from the Northern Netherlands population cohort Lifelines. For each individual, peripheral blood mononuclear cells (PBMCs) were sequenced in an unstimulated condition and after 3h and 24h *in vitro*-stimulation with *C. albicans* (CA), *M. tuberculosis* (MTB) or *P. aeruginosa* (PA), totaling approximately 1.3 million cells. First, we identified the cell type⍰specific transcriptional changes in response to these pathogen stimulations. Subsequently, we determined how genetics may affect these gene expression responses. This resulted in a final set of genes that were differentially expressed and showed cell type⍰specific and/or stimulation-dependent eQTL effects. For this final gene set, we performed co-expression QTL analysis to test if the associated eQTL SNPs also affect co-expression between the eQTL gene and other genes. Interestingly, this analysis showed many cases of pathogen- and/or time-dependent genetic control of gene regulation, revealing novel avenues through which genetics can contribute to disease risk.

## Results

### Single-cell profiling of immune cells upon pathogen stimulation

We performed 10x Genomics scRNA-seq on PBMCs from 120 individuals that were stimulated with three different pathogens at two timepoints, or left unstimulated (**Fig. 1, Table S1**). A combination of 10X Genomics v2 and v3 chemistry reagents were used to capture an average of 1,226 cells per individual per condition (v2: 907 genes/cell, v3: 1,861 genes/cell). Souporcell^29^ was used to identify the doublets coming from different individuals, followed by sample demultiplexing using Demuxlet.^11^ This revealed, on average, 12.0% of cells as doublets. Due to differences in gene amplification between v2 and v3 chemistry, determination of quality control (QC) thresholds and analyses were performed separately per chemistry (**Fig. S1a**). Results from both chemistries were then meta-analyzed for interpretation. Low quality cells were excluded, leaving 928,275 cells in the final dataset used for analyses (see Methods). UMAP dimensionality reduction and KNN-clustering was then applied on the normalized, integrated count data, allowing the identification of six main cell types: B, CD4+ T, CD8+ T, monocytes, natural killer (NK) and dendritic cells (DCs) (**Fig. S1b-d**).

**Figure 1.**
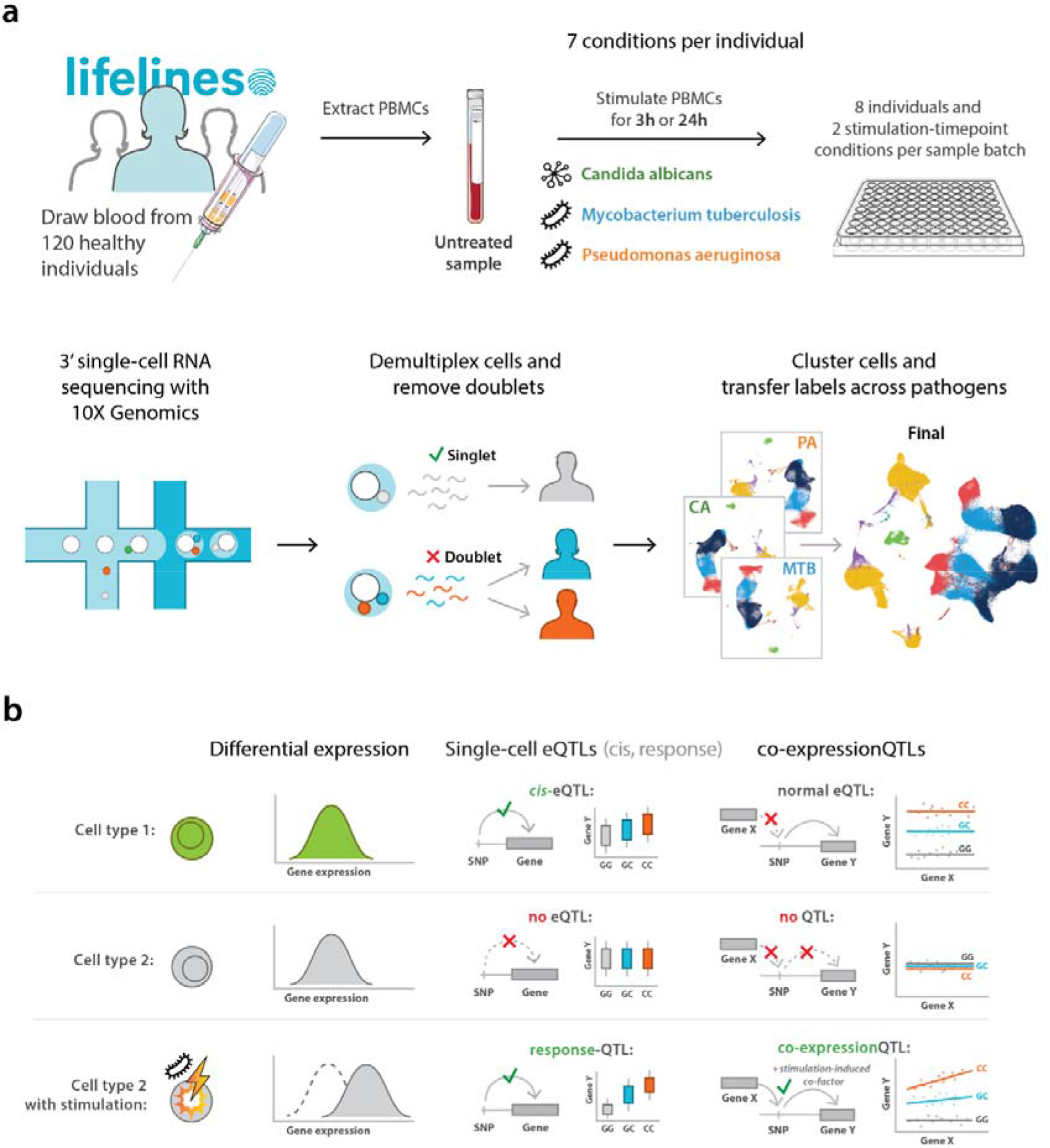
Study overview. **a**. Blood was drawn from 120 individuals from the Northern Netherlands participating in the population-based cohort Lifelines. PBMCs were isolated within 2h of blood collection and cryopreserved in liquid nitrogen until further use. For each scRNA-seq experiment, PBMCs from 16 individuals were thawed. Per individual, these PBMCs were left untreated (UT) or were stimulated with C. Albicans (CA), M. Tuberculosis (MTB) or P. Aeruginosa (PA) for 3h or 24h in a 96-well plate. In total, this resulted in 7 stimulation⍰timepoint combinations per 120 individuals = 840 different conditions processed. Multiplexed 3’-end scRNA-seq was performed using the v2 and v3 chemistries of the 10X Genomics platform. Per experiment, two sample batches of a 10x chip were loaded, each containing a mixture of eight individuals and a combination of two different stimulation⍰timepoint combinations. After sequencing, samples were demultiplexed and doublets were identified. Cell type classification was performed on the QCed dataset by clustering the cells per pathogen, mixed with cells of the unstimulated condition. The cell type labels were subsequently transferred back to the dataset containing all cells. **b**. This study was conducted to identify cell type⍰specific or pathogen-stimulation dependent: 1. differentially expressed genes, 2. eQTLs and response-QTLs and 3. co-expression QTLs.

### Gene expression response upon pathogen stimulation reveals stronger cell type⍰specificity than pathogen-specificity

To assess the transcriptional changes upon pathogen stimulation with CA, MTB and PA, we performed differential expression (DE) analysis using MAST (**Table S2**).^30^ Pairwise comparisons between the untreated and pathogen-stimulated conditions revealed between 688 to 2,022 DE genes after 3h stimulation, further increasing to 1,052 to 2,616 DE genes after 24h stimulation (**Fig. 2a**). The number of DE genes was comparable between the different pathogen stimulations at the same timepoint but differed strongly between some cell types. Myeloid cells (monocytes and DCs) showed the highest number of DE genes, whereas both CD4+ and CD8+ T cells showed the fewest DE genes. This is consistent with the innate immune cells being the first responders during pathogen stimulation.^31^

**Figure 2.**
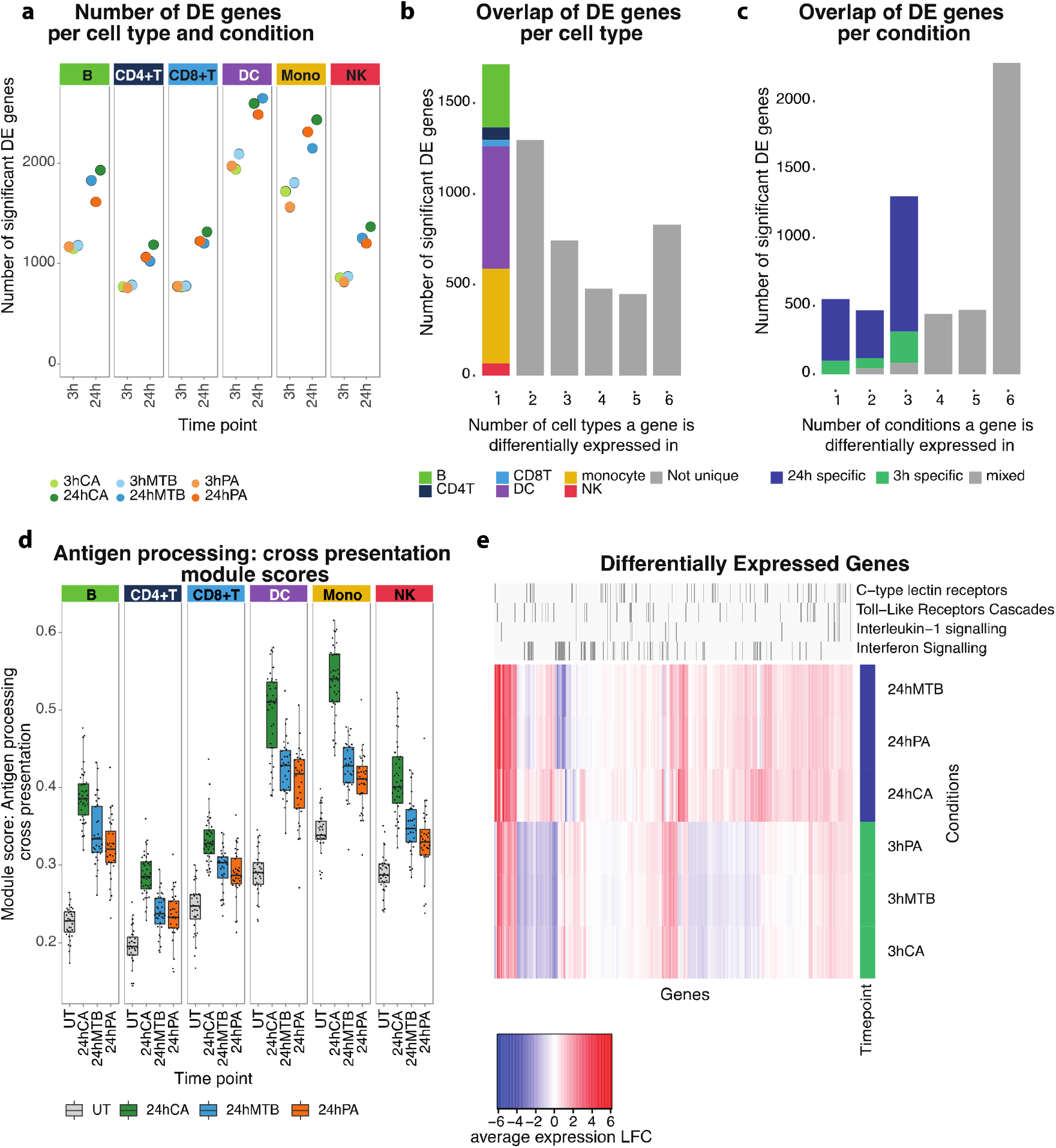
Differentially expressed genes and pathways upon pathogen stimulation. **a**. Number of DE genes per cell type upon 3h (light colors) or 24h (dark colors) stimulation with C. Albicans (CA, green), M. Tuberculosis (MTB, blue) or P. Aeruginosa (PA, orange). **b**. Bar plot showing the overlap of DE genes across cell types. The first bar, depicting the cell type-specific DE genes, is colored based on the cell type in which the DE gene is found. **c**. Bar plot showing the overlap of DE genes across pathogen⍰timepoint combinations (3h vs 24h stimulation with CA, PA or MTB). Bars are colored based on the length of stimulation. **d**. Boxplots (showing median, 25 ^th^ and 75^th^ percentile, and 1.5 ×flthe interquartile range) representing the module score of the antigen-processing cross-presentation pathway across all individuals per cell type and per 24 hours pathogen-stimulated condition. Each dot shows the average module score per individual. **e**. Heatmap showing the immune-related DE genes in monocytes (V3 chemistry is shown). The DE genes in four immune pathways that are highlighted on the top are the main pathways involved in pathogen recognition and its downstream signaling. C-type lectin and toll-like receptors show more general activation upon pathogen stimulation, whereas interleukin-1 and interferon signaling show a more specific expression pattern with timepoint (3h stimulation) or stimulation (CA), respectively.

A total of 2,643 unique DE genes were identified over all conditions and cell types. Of these, 31.1% were cell type⍰specific and 15.1% were shared across all cell types (**Fig. 2b**). The fraction of DE genes that were cell type⍰specific was comparable for each of the cell types, but, in absolute numbers, monocytes and DCs had the most unique DE genes. Sharing between different pathogen stimulations at the same timepoint was more prominent than sharing between different timepoints within the same pathogen stimulation (**Fig. 2c, Fig. S1c**): 39.8% of the total unique DE genes were shared across the same timepoint (7.4% at 3h and 32.4% at 24h), whereas only 10.3% of DE genes were unique to a specific pathogen stimulation and 41.3% were shared across all stimulation⍰timepoint combinations. This indicates that the immune response to our pathogen stimulations of both bacterial and fungal origin was more specific to timepoint after stimulation than to type of pathogen. Consequently, the genetic control of these responsive genes is expected to be more time-dependent than pathogen-dependent.

To evaluate the DE results and confirm proper activation of the cells upon stimulation, we performed two different analyses. In the first analysis, we measured the activity of a general stimulation-responsive pathway ⍰ the antigen processing-cross presentation pathway (REACTOME R-HSA-1236975) ⍰ that should become activated in each of the cell types and upon each of the pathogen-stimulations. This analysis revealed increased activity of the antigen-processing pathway-associated genes across all cell types after 24h stimulation and for each of the pathogens (**Fig. 2d**). In the second analysis, we focused on DE genes identified upon 24h stimulation with CA. We had previously performed similar analyses in a smaller scRNA-seq study^15^, so we could use this study for comparison purposes. This analysis revealed a high concordance between DE genes in our current study and those from our previous study, varying from 73% for the monocytes up to 93% for the B cells (**Fig. S2**). In general, these analyses showed that CA stimulation resulted in the highest activation of genes associated with the antigen-processing pathway and that monocytes were the cell type with the strongest response. These two analyses confirmed proper activation of the cells and stimulation responses that were in line with previous literature.^15,32^

Next, we determined which pathways were enriched within the upregulated DE genes for each cell type and each pathogen⍰timepoint combination (**Table S3**). In line with the DE results, most of the enriched pathways were shared across the different pathogen stimulation conditions within the same timepoint (**Fig. 2e**). To highlight relevant pathways involved in pathogen recognition and downstream immune response, we filtered the enriched pathways for those related to the ‘Immune system’ REACTOME pathway parent term. For this illustrative example, we selected monocytes because this was the cell type in which we observed the most DE genes (**Fig. 2e**). Here we observed a general activation of pathogen-recognition receptors and downstream signaling, including the C-type lectin and toll-like receptors. Some pathways, such as interleukin-1 (IL-1) signaling, were clearly enriched at a specific timepoint (3h stimulation), whereas others, such as the IFN pathway, showed a notable difference between different pathogen stimulations (more prominently activated in CA compared to the other two pathogens). These findings corroborate literature describing IFN as an important signaling pathway in response to all three pathogens^32–34^ and that IL-1 family molecules are part of the early stages (<14h) of the inflammatory response in monocytes with their expression decreasing again at later stages.^35^

### The number of eQTLs decrease in cells with stronger stimulation response

Our experimental set-up, in which we analyzed pathogen-stimulated PBMCs using scRNA-seq, allowed us to investigate the extent to which SNPs affect gene expression in different contexts. To maximize the power to detect eQTLs, we took advantage of a previously conducted genome-wide *cis*-eQTL meta-analysis in 31,684 whole blood bulk samples (eQTLGen^36^) by only testing their top SNP⍰gene combinations, i.e. lead-eSNPs. Due to the power of eQTLGen, it could identify even *cis*-eQTL effects with a small effect size. We therefore expected that many of the context-specific effects, to which only a subset of individuals or cell types might have been exposed, should have resulted in an eQTL effect identified in eQTLGen. However, compared to the eQTLGen bulk whole-blood dataset, our pathogen-stimulated scRNA-seq data has the additional benefit that it can identify the cell types and contexts in which these eQTL effects manifest themselves.

We performed the eQTLGen lead-eSNP *cis*-eQTL discovery analysis per cell type and for each stimulation⍰timepoint combination separately (**Table S4**). When determining the concordance between eQTLGen’s bulk whole-blood eQTLs and those identified in our study, we observed that the concordance was high in general despite the compositional differences between whole blood and the PBMCs or cells in this study that were pathogen-stimulated. As expected, we obtained the highest concordance (95.5%) with eQTLGen when comparing to our bulk-like unstimulated PBMC scRNA-seq data, i.e. taking the average gene expression across all cells from one individual in the untreated condition (**Fig. 3a**). We then saw only a minor drop to 94.7% concordance when comparing eQTLs from eQTLGen with our pathogen-stimulated (24h CA) bulk-like scRNA-seq dataset (**Fig. 3b**) and a further decrease to 92.6% when comparing to our pathogen-stimulated and cell-type⍰specific scRNA-seq dataset (24h CA in monocytes) (**Fig. 3c**). Finally, to verify that our initial selection of eQTLGen lead-eSNPs did not confound our conclusions, we also compared the output of a genome-wide *cis*-eQTL discovery (**Table S5**) in pathogen-stimulated and cell type⍰specific scRNA-seq data (24h CA in monocytes) with eQTLGen. In this analysis, the concordance decreased a little bit further to 87.4% (**Fig. 3d**). Although up to 19.6% of the eQTLs were only detected in the genome-wide discovery (and not in the eQTLGen lead-eSNP-confined *cis*-eQTL discovery), these unique *cis*-eQTL gene sets were not enriched for specific biological pathways. Altogether, this indicated that the eQTLGen lead-eSNP confinement only had a minimal impact on our observations and confirmed our initial assumption that the majority of context-specific eQTLs identified by our current study can already be detected in very large bulk RNA-seq datasets. However, we still require single-cell data to pinpoint their relevant context. As the eQTLGen lead-eSNP *cis*-eQTL analysis identified 1.5x more eQTLs, while showing no clear bias towards common eQTLs rather than cell type⍰specific or context-dependent eQTLs (*Fig. S3a*), we continued our analysis with these results.

**Figure 3.**
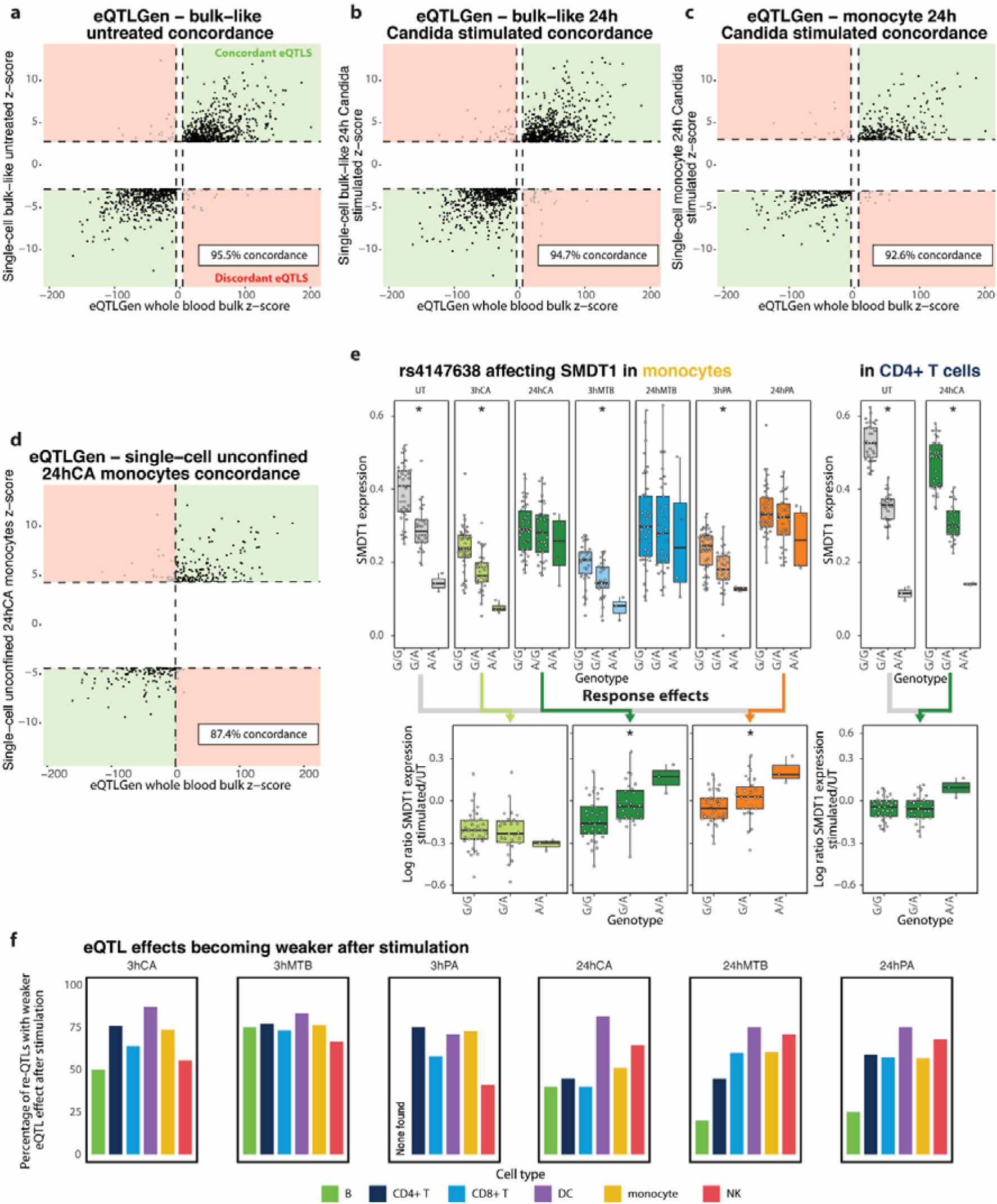
eQTLs and re-QTLs upon pathogen stimulation. Concordance between the eQTLs identified in 31,684 bulk whole blood samples of the eQTLGen consortium and: **a**. those identified in our eQTLGen lead-eSNP discovery of bulk-like unstimulated PBMC scRNA-seq data, **b**. those identified in our eQTLGen lead-eSNP discovery of bulk-like 24h C. Albicans (CA)-stimulated PBMC scRNA-seq data, **c**. those identified in our eQTLGen lead-eSNP discovery of monocyte 24h CA-stimulated PBMC scRNA-seq data, and **d**. those identified in our genome-wide eQTL discovery of monocyte 24h CA-stimulated PBMC scRNA-seq data. **e**. Box plots showing the effect of the rs4147638 genotype on SMDT1 expression in the untreated (UT) condition and each of the six stimulationfltimepoint combinations in the monocytes (left) or for the UT and 24h CA condition in the CD4+ T cells (right). Box plots show median, first and third quartiles, and 1.5fl×flthe interquartile range, and each dot represents the average expression of all cells per cell type and individual. Stars indicate a significant effect (FDR < 0.001). The log ratio of SMDT1 expression in the UT cells vs a specific stimulation-timepoint combination is shown in the bottom. Colored arrows indicate which specific stimulationfltimepoint combination was selected for the corresponding re-QTL boxplot. **f**. The proportion of re-QTLs of which the eQTL effect became weaker after stimulation, split per cell type and stimulationfltimepoint combination.

The CD4+ T cells revealed the most eQTL effects, followed by the monocytes and CD8+ T cells (**Fig. S3a**). The cell types with the lowest frequencies, the DCs and B cells, also showed the lowest number of eQTLs (**Fig. S3b**). This large difference in the number of identified eQTLs per cell type is, at least in part, explained by the differenced in power, given the number of cells of each cell type (**Fig. S1d**). In addition to differences between cell types, we also observed differences between stimulation⍰timepoint combinations (**Fig. S3c**). However, direct comparisons of the number of eQTLs between conditions within the same cell type were complicated because the number of included individuals varied among the stimulation⍰timepoint combinations as a result of QC dropouts (UT: 104 individuals, 3h CA: 120 individuals, 3h MTB: 104 individuals, 3h PA: 112 individuals, 24h CA: 119 individuals, 24h MTB: 112 individuals, 24h PA: 111 individuals). Most interestingly, when comparing the effect of pathogen stimulation on the number of identified eQTLs between cell types, we observed an inverse correlation with the responsiveness of that cell type to pathogen stimulation (**Fig. S3d**). For example, the myeloid cells showed the largest DE response upon pathogen stimulation (**Fig. 2a**) but a consistent reduction in the number of eQTLs identified after stimulation (**Fig. S3a**). In contrast, the lymphoid cells showed a much smaller DE response upon pathogen stimulation (**Fig. 2a**) but an increase in the number of eQTLs identified after stimulation, in about half of the conditions (**Fig. S3a**). This could indicate that, for at least a subset of the genes, the influence of genetics on gene expression may become more restricted when cells have to orchestrate a response to an environmental stimulus.^37^

To identify eQTLs for which the strength of the eQTL effect was affected by pathogen stimulation, we performed a response-QTL (re-QTL) analysis.^38^ We systematically looked for re-QTLs in all cell types and stimulation conditions compared to the untreated condition (**Table S6**). Most re-QTLs were specific to a particular timepoint or cell type, but less so to a particular pathogen (**Fig. 3e**). We observed that most re-QTLs were in the monocytes for each of the stimulation⍰timepoint combinations (**Fig. S3a**), likely the direct consequence of the combination of a high number of DE genes upon stimulation (**Fig. 2a**) and the relatively high number of monocytes per individual (**Fig. S1d**). We also observed that most re-QTLs describe eQTL effects that became weaker after stimulation (**Fig. 3f**). Of those eQTL effects that became stronger after stimulation, 26.3% on average showed a significant effect that was already present in the unstimulated samples, whereas those effects were not yet present for 63.7%. Moreover, we observed clear enrichment of DE genes within the set of eQTL and re-QTL genes, but this enrichment was not consistently greater for re-QTL in comparison to eQTL genes (**Fig. S3e**).

Finally, when linking the eSNP loci to GWAS output of immune-mediated diseases (see Methods), we observed a strong genomic inflation across all conditions (**Table S7**). This genomic inflation increased further for the re-QTLs (in monocytes over all immune-mediated GWASes: p=0.024) (**Table S7**). These findings confirmed previous studies showing that stimulation-responsive eQTL effects provide additional explanation of immune-mediated disease risk over baseline eQTLs^4,39^. Additionally, it has been shown that the effect size of GWAS-associated SNPs becomes larger in the disease-relevant context (e.g. immune-mediated disease patients as opposed to the healthy controls).^40^ Therefore, also the power to detect these disease-associated effects will be larger in the disease relevant context.

In summary, we observed that 20.9% of our eQTL genes were influenced by a combination of genetics and environment (**Table S4, S6**). We expect this percentage is an underestimate, as the power to detect re-QTLs is inherently lower than that of eQTLs and exposure to additional environmental stimuli may reveal additional context-dependency. Altogether, our findings indicate that, in addition to cell type⍰specificity, context-dependency is also a major driver of genetic regulation of gene expression and provides additional explanation of disease risk.

### Pathogen stimulation induces widespread context-specific gene regulation

We have previously shown that genetics can influence the co-expression relationship between genes.^12^ Similarly, studies that compared co-expression in healthy versus disease states have indicated that environmental conditions may also impact these gene⍰gene interactions.^41^ Here, we took the next step by determining whether and how the combination of genetics and environment may affect how genes are interacting with one another by performing co-expression QTL analysis, i.e. a SNP genotype affecting the co-expression relationship of a gene pair. For this purpose, we selected a subset of 49 SNP⍰gene combinations that we then tested against up to 5,772 genes. To enrich for SNP⍰gene combinations in which we expect an interaction with the environment, we selected these based on: 1. the gene being DE and 2. the SNP⍰gene combination being a re-QTL in at least one of the stimulation⍰timepoint combinations; 3. the gene being expressed in at least 50% of the individuals (in each 10X chemistry). For this analysis, we focused solely on the monocytes because this was the cell type that showed a strong response to pathogen stimulations and for which we had sufficient cells per individual (i.e. hundreds) to perform a robust co-expression QTL analysis. By making this pre-selection of 49 SNP⍰gene combinations, we could reduce the multiple testing burden from over 10^14^ in a genome-wide analysis to fewer than 283,000 tests.

Across the unstimulated condition and each of the six stimulation⍰timepoint combinations, we found at least one co-expression QTL for 35 SNP⍰gene combinations and more than 100 co-expression QTLs in at least one condition for 9 SNP⍰gene combinations. For each of these 9 SNP⍰gene combinations with a high number of co-expression QTLs, we observed an interaction between genotype and stimulation condition (**Fig. 4a, Table S8**). One of these co-expression QTLs described an interaction between RPS26 and rs1131017, which was an effect in high LD with one we had identified as a co-expression QTL in CD4+ T cells in our previous study (rs7297175, R^2^ = 0.92). ^12^ rs1131017 was previously associated with rheumatoid arthritis^42^ (p = 1.3×10^−8^) and is in high LD with a type I diabetes GWAS SNP^43^ (rs11171739, R^2^ = 0.94). For this *RPS26*⍰rs1131017 SNP combination, we found 1,701 co-expression QTLs in the unstimulated condition. Of the 106 *RPS26* co-expression QTLs that we had previously identified in CD4+ T cells^12^, 72 (67.9%) were also found in the unstimulated monocytes in our current study (91.7% with the same allelic direction) (**Fig. S4a**). Any discrepancy between these two cell types might be the consequence of distinct regulatory mechanisms that are active in those cell types. Next, looking at the effect of stimulation, the number and strength of the detected *RPS26* co-expression QTLs reduced greatly after stimulation and was related to the duration of stimulation: on average we observed 459 co-expression QTLs after 3h stimulation and 112 after 24h stimulation (**Fig. 4a, Fig. S4b**).

**Figure 4.**
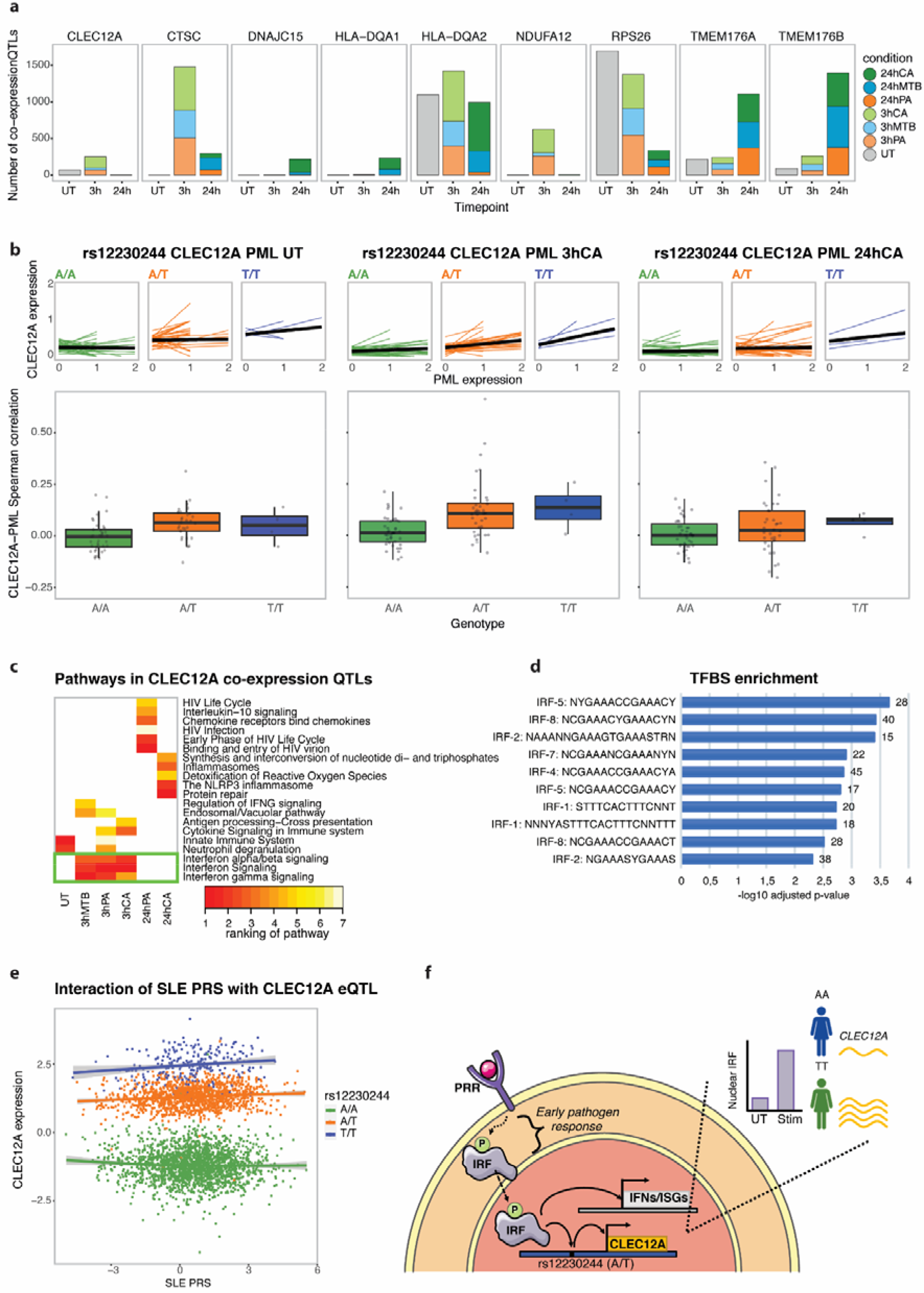
Interferon regulatory factor affects CLEC12A co-expression QTLs upon 3h pathogen stimulation in monocytes. **a**. Number of co-expression QTLs identified in each of the stimulationfZtimepoint combinations for those co-expression QTLs with over 100 co-expression QTLs in at least one condition. The 3h and 24h timepoint are colored by pathogen stimulation (green: C. Albicans (CA), blue: M. Tuberculosis (MTB), orange: P. Aeruginosa (PA). **b**. The lines in the top plots show co-expression between CLEC12A and PML (most significant co-expression QTL across the 3h stimulation conditions) for individual cells in the untreated (left), 3h CA (middle) and 24h CA (right) condition. In these plots, individual-specific regression lines are shown, split by genotype. The average genotype-specific regression lines are shown in black. The bottom boxplots depict Spearman’s rank correlation between CLEC12A and PML expression, stratified by SNP rs12230244 genotype in the monocytes per individual, in the untreated (left), 3h CA (middle) and 24h CA (right) stimulated cells (the V2 chemistry data is plotted). Each data point shows a single individual. Box plots show median, first and third quartiles, and 1.5× the interquartile range. **c**. Heatmap of the top-5 enriched pathways within the co-expressed CLEC12A co-eQTL genes per stimulationfZtimepoint combination. Per combination, pathways are ranked based on significance. White indicates that the pathway was not found to be enriched in that specific stimulationfZtimepoint combination. The green box highlights pathways that are associated with all 3h stimulation conditions. **d**. Top 10 enriched putative transcription factor binding sites within the CLEC12A co-expression QTL genes that: 1. showed a more positive strength of the co-expression relationship in individuals with the TT as opposed to the AA genotype and 2. were identified in the 3h stimulated (outer join) monocytes using the TRANSFAC database. **e**. Co-expression QTL analysis for CLEC12A-SNP rs12230244 against the SLE PRS (calculated using those SLE GWAS SNPs with a P-value threshold of <5×10 ^-8^) using whole blood bulk expression data from 3,553 individuals (BIOS consortium). **f**. Proposed mechanism of action of CLEC12A co-expression QTLs. When pathogen-associated molecular patterns bind to a pattern recognition receptor (PRR), a signaling cascade is initiated that eventually results in phosphorylation of interferon regulatory factors (IRFs). Phosphorylated IRF then translocates to the nucleus, where it binds to specific DNA motifs such as IFN-stimulated response elements. This can then activate transcription of IFNs and IFN-stimulated genes (ISGs). Additionally, IRF is expected to bind to a region containing SNP rs12230244 (or any another SNP in high LD), thereby regulating CLEC12A expression. In this case, depending on the SNP genotype, the IRF binding and activation of CLEC12A expression is expected to be stronger (TT genotype) or weaker (AA genotype). Many of the identified CLEC12A co-expression QTL genes are involved in the IFN pathway (see **4b**). This has to be the result of a common upstream factor (i.e. IRF) of CLEC12A transcription that can also activate IFNs and ISGs, but cannot be the result of a downstream regulator because this would have led to trans-eQTL effects for the same SNP rs12230244 (which we do not observe).

We also observed this general decrease in the strength and number of co-expression QTLs with increasing duration of pathogen stimulation for the *HLA-DQA2* co-expression QTLs, but not for any of the other 7 SNP⍰gene combinations (**Fig. 4a**). These other 7 co-expression QTL effects increased in strength and numbers upon stimulation (**Fig. 4a, Table S8**). Interestingly, for some of these co-expression QTL genes, we observed the most prominent increase at 3h stimulation (i.e. *CLEC12A, CTSC* and *NDUFA12*), whereas others were more prominent at 24h stimulation (i.e. *TMEM176A*/*B, DNAJC15* and *HLADQA1*). The observation of different numbers of co-expression QTLs for a specific gene over the 7 stimulation⍰timepoint combinations was not fully explained by the expression level of that gene. Beyond this variation over the timepoints, we also observed clear differences between the various pathogen stimulations. At gene-level, there was little overlap between the co-expressed gene sets between the different pathogen stimulations (**Table S8**), whereas this overlap was much larger at pathway-level (**Fig. S5**). The low gene-level overlap is likely a consequence of power and is something that will be largely overcome in the near future with the increase in the number of cells per dataset.^44,45^ Together, these results indicate that specific environmental conditions can fulfill the requirements needed for a specific co-expression QTL interaction to occur.

Previously, re-QTL analyses in cells exposed to highly specific stimuli were used to disentangle the environmental conditions that underlie specific genetic regulation of gene expression.^4,16^ However, this has the disadvantage that either many highly specific stimuli have to be applied, or, in the case of applying broad stimuli, the exact environmental context relevant for the interaction remains vague. Here, we propose using co-expression QTL analysis upon stimulation with a few broad stimuli to gain this detailed insight in a more unbiased way, without the need to apply many highly specific stimuli. As a first example of how co-expression QTL analysis can help us understand the underlying mechanisms of gene regulation, we focused on the *CLEC12A* co-expression QTLs affected by SNP rs12230244, which were most prominent at 3h of pathogen stimulation (**Fig. 4a, 4b**). *CLEC12A*, also known as *MICL*, encodes for an inhibitory C-type lectin-like receptor and is mostly expressed in myeloid cells such as monocytes and DCs. CLEC12A signaling can be activated by the binding of uric acid crystals, which are the byproduct of nucleic acids that can be released from damaged or dying cells.^46,47^ Activation of CLEC12A signaling can result in inhibition of the activating C-type lectin receptors and can prevent hyperinflammation during necrosis.^48^

To identify the potential causal factor underlying the *CLEC12A* co-expression QTL, we performed a pathway analysis on the associated co-expressed gene set of each of the stimulation⍰timepoint combinations. We hypothesized that co-expressed genes linked to the same co-expression QTL mostly describe the same (or only a few) biological processes that are driven by a single (or a few) causal factors being directly involved, and that most of these co-expressed genes are themselves just a consequence of being highly co-expressed with the causal factor. An important category of causal factors are transcription factors. However, average expression levels of transcription factors are generally low and, particularly in dynamic situations such as a pathogen response, mRNA levels might not correlate well with the nuclear protein expression levels (i.e. the functional proportion).^49,50^ Consequently, it can be difficult to define the direct causal factor solely using co-expression QTL analysis. Nevertheless, we expected that by taking a pathway-level view, the downstream genes of transcription factors would have a high correlation with the functional protein level of the transcription factor and would be more easily picked up than a single gene.

The pathway analysis of the *CLEC12A* genotype-dependent co-expressed gene set after 3h stimulation (Benjamini-Hochberg (BH) corrected p = 2.9×10^−5^, 8.7×10^−7^ and 4.3×10^−4^ for 3h CA, 3h PA and 3h MTB, respectively), but not in the untreated or 24h stimulation conditions, revealed enrichment of the IFN pathway (**Fig 4c**). This result hinted that a component within or regulating the IFN pathway could be the causal factor that is regulating the different *CLEC12A* co-expression responses per genotype after 3h stimulation. To provide additional support for this hypothesis, we performed a functional enrichment analysis for putative transcription factor binding sites (TFBSs) (TRANSFAC database^51^) on the *CLEC12A* genotype-dependent co-expressed genes upon 3h stimulation. We divided this gene set into a subset in which individuals with the TT as opposed to the AA genotype showed a more positive rather than a more negative co-expression relationship between *CLEC12A* and its co-expressed genes, as potentially different mechanisms could be underlying these gene sets. This analysis revealed no enriched TFBSs in the negative-strength gene set, but a clear enrichment of various IFN regulatory factors (IRFs) in the positive-strength gene set, including *IRF1, 2, 4, 5, 7* and *8* (**Fig. 4d**). Additionally, when overlapping the *CLEC12A* co-expression QTL SNP rs12230244 and its accompanying (near-)perfect LD SNPs with putative TFBSs^52^, we observed several transcription factors that may bind to the genomic location of these SNPs. Most noticeably, the predicted binding site of *IRF1* was shown to be enriched in the genomic location of two SNPs that are in near-perfect LD with the *CLEC12A* co-expression QTL SNP: rs999185 (R^2^ = 0.9943) and rs57106602 (R^2^ = 0.9).

Finally, we used two external datasets and slightly different approaches to further strengthen our hypothesis that IFN activity is regulating the *CLEC12A* co-expression QTL effects. First, we used the BIOS consortium bulk RNA-seq dataset containing whole blood data from 3,553 individuals.^36^ For each of those individuals, we calculated a polygenic risk score (PRS) for the autoimmune disease systemic lupus erythematosus (SLE), a disease characterized by increased type I IFN activity.^53–56^ We reasoned that the genetic risk captured by the SLE PRS could be used as a proxy for IFN activity. Consequently, the difference in the co-expression relationship between the SLE PRS and the *CLEC12A* per rs12230244 genotype indicated the involvement of IFN signaling in this interaction (**Fig. 4e**). Second, we used an independent scRNA-seq dataset generated in 68 healthy controls and 117 SLE patients from European (EUR) and East Asian (EAS) origin. We reasoned that since IFN activity is characteristic for SLE^53–56^, SLE patients would mimic the 3h pathogen stimulation state in which high IFN activity seems to drive the observed *CLEC12A* co-expression QTL effects. We also reasoned that the healthy controls would mimic the untreated cells in our study and therefore show fewer *CLEC12A* co-expression QTL effects driven by IFN activity. To define whether the SLE patients mimicked the results as observed after 3h pathogen stimulation, we performed a co-expression QTL analysis for *CLEC12A* and SNP rs12230244 in the monocytes of SLE patients and healthy controls (*Table S9*). Pathway analysis on the *CLEC12A* co-expression QTL genes revealed stronger enrichment for the IFN pathway in the SLE patients (FDR = 1.965×10^−7^) compared to the healthy controls (FDR = 1.203×10^−3^), again supporting that this pathway is involved in the regulation of *CLEC12A* through SNP rs12230244.

As a second example, we applied a similar strategy to learn the underlying regulatory mechanism by which the co-expression QTLs identified for SNP rs6945636 affect the heat-shock protein response gene *ZFAND2A*. The heat-shock protein response is a pathway that, amongst others, can be activated by bacterial and viral infections.^57^ We selected this specific SNPCgene combination for further analysis because the co-expression QTLs identified were both pathogen- and timepoint-specific (only at the 24h timepoint, 96% of the genes being detected in CA only). Pathway analysis of the co-expressed genes revealed ‘Intracellular pH reduction’ (GO:0051452) as the top associated biological process (adjusted p = 3.8×10^−4^). Interestingly, *HSF1*, a known regulator of *ZFAND2A*^58^ was shown to be pH sensitive in yeast.^59^ Moreover, the *ZFAND2A*-associated co-expression QTL SNP rs6945636 was in almost perfect LD with previously identified *HSF1* binding sites in K562 cells (R^2^ = 0.99, rs715188378; R^2^ = 0.99, rs79849558; R^2^ = 0.99, rs11767061, retrieved from dbSNP release 153).^60^ Together, this indicates that CA-induced pH regulation activated *HSF1*, which in turn bound with stronger (TT genotype) or weaker (AA genotype) strength to SNP rs6945636 (or any other in high LD) and thereby strongly or weakly activated *ZFAND2A*, respectively.

These two examples provide clear use cases for how co-expression QTL analysis can be applied to gain detailed insights into the underlying context of gene expression regulation. For example, in the case of *CLEC12A*, without co-expression QTL analysis we could only reveal that *CLEC12A* is a re-QTL regulated by a factor active 3h downstream of pathogen stimulation (*Table S6*). In contrast, using co-expression QTL analysis, we were directed to the causal regulatory factor for this re-QTL. This enabled follow-up analyses that gathered solid evidence for the following mechanism of action through which SNP rs12230244 affects *CLEC12A* expression specifically upon 3h pathogen stimulation: 1. pathogen-associated molecular patterns bind to a pattern recognition receptor (PRR) and initiate a signaling cascade that eventually results in phosphorylation of interferon regulatory factors (IRFs), 2. phosphorylated IRF then translocates to the nucleus where it binds to specific DNA motifs such as IFN-stimulated response elements, and 3. this can then activate transcription of IFNs and IFN-stimulated genes (ISGs). Additionally, IRF is expected to bind to a region containing SNP rs12230244 (or any another SNP in high LD), thereby regulating *CLEC12A* expression. In this case, depending on the SNP genotype, the IRF binding and induction of *CLEC12A* expression is expected to be stronger (TT genotype) or weaker (AA genotype) (**Fig. 4f**).

Interestingly, we identified a number of (near-)genome-wide significant PheWAS traits related to immune cell composition and size to be associated with these two co-expression QTL loci (extracted from 452,264 White British individuals of the UK Biobank^61^): platelet counts (p = 2.1×10^−8^), monocyte percentage (p = 1.8×10^−5^) and eosinophil counts (p = 6.4×10^−5^) for *CLEC12A* and mean corpuscular volume (p = 1.5×10^−14^) and mean sphered cell volume (p = 2.6×10^−9^) for *ZFAND2A*. However, no direct association was found for any of the immune-related GWAS tested (SLE^62^, inflammatory bowel disease^63^, celiac disease^64^, rheumatoid arthritis^42^, multiple sclerosis^65^, type I diabetes mellitus^66^ and candidemia^67^, which had 10-2,541-fold smaller sample sizes than the PheWAS. This overlap with immune-related PheWAS traits indicate the relevance of these SNPs for immune function. Using co-expression QTL analysis, we can now dissect the underlying mechanism by which such an effect is regulated. This information will help explain the downstream consequences on immune function, and potentially enable new routes for medical intervention.

## Discussion and Conclusion

GWAS studies have provided important insights into the genetic architecture of phenotypic traits and diseases.^1^ However, the exact mechanisms by which genetic variation leads to these traits or diseases largely remain a black box. To uncover these mechanisms, various approaches have been successfully applied, for example coupling the trait-associated risk factor to the nearest positional gene, downstream gene expression^36^, or gene regulation.^12^ Nevertheless, a large knowledge gap remains that may, in part, be filled by taking into consideration the context in which the genetic variant can lead to disease.^7,16,17^

To uncover the interplay between genetics and cellular and environmental context, we single-cell RNA-sequenced PBMCs from 120 individuals from Lifelines, a large population-based cohort from the Northern Netherlands, that had been exposed to various pathogens or left untreated. Subsequent DE, eQTL and co-expression QTL analysis revealed that there are widespread interactions between an individual’s genetics and the cellular and environmental context, both at the level of gene expression and in its regulation. We identified hundreds of eQTLs in the individual cell types and upon pathogen-stimulation and observed strong context-specificity for 25.7% of the identified co-expression QTLs. In general, we observe more interactions between genetics and cell typeCspecific context, as opposed to context induced by pathogen stimulation. However, some of these differences may have been the result of differences in detection power. Contrary to expectations, in the cell types with the strongest response to pathogen stimulation (i.e. the myeloid cells), the total number of eQTLs was reduced after stimulation. Moreover, in all cell types, we observed that eQTLs more often became weaker rather than stronger after stimulation and that neither category of eQTL genes was associated with a specific pathway. In contrast, for the co-expression QTL genes, the number of co-expressed genes more often increased upon pathogen stimulation. However, this might in part have been the result of our selection, i.e. choosing re-QTLs in monocytes as the starting point for the co-expression QTL mapping. Moreover, we observed genomic inflation of eQTLs that further increased when focusing solely on the re-QTLs. Altogether, these observations indicate that context, here the pathogen-stimulation condition, is an important contributor that affects the association between SNPs and gene expression or co-expression, and that taking this context in consideration further improves our understanding of disease risk.

A major advantage of co-expression QTL analysis as opposed to re-QTL analysis is that we do not require many highly specific stimuli to disentangle the mechanisms that underlie the context-specificity of the genetic regulation. Instead, in this study, we have shown that, after applying a broad stimulation (i.e. whole-pathogen stimulation), a wide range of contexts are activated, and that, through subsequent co-expression QTL analysis, the specific context and mechanism of action could be uncovered. For example, we revealed that an interferon-regulated transcription factor was affecting the SNP rs12230244⍰dependent downstream activation of *CLEC12A*. Additionally, we showed how pH-dependent regulation of the heat shock protein response transcription factor *HSF1* affected the SNP rs6945636⍰dependent downstream activation of *ZFAND2A*. These examples clearly show the potential of the technology and provide an outlook into where the field will be moving as more population-scale scRNA-seq datasets become available.

In the last few years, scRNA-seq has become a mature, high-throughput technology.^8,9^ This has led to several initiatives aiming to study population genetics at single-cell resolution, such as the sc-eQTLGen consortium^44^ and others.^68^ Such efforts bring together many single-cell eQTL studies, conducted in individuals from different ethnicities and exposed to different environments or diseases. This will not only increase the power to detect eQTLs and co-expression QTLs, it will also further extend our findings to additional contexts and enable genome-wide cell-type and context-specific *trans*-eQTL mapping. By integrating GWAS signals, PRS scores and context-specific QTL information, we expect that these efforts can drive major leaps forward in disease understanding and precision medicine.^69^

## Materials and methods

### PBMC collection and stimulations

Whole blood from 120 individuals of the northern Netherlands population cohort Lifelines Deep^70^ was drawn into EDTA-vacutainers (BD). PBMCs were isolated and maintained, as previously described.^12^ In short, PBMCs were isolated using Cell Preparation Tubes with sodium heparin (BD) and were cryopreserved until use in RPMI1640 containing 40% FCS and 10% DMSO. After thawing and a 1h resting period, unstimulated cells were washed twice in medium supplemented with 0.04% BSA and directly processed for scRNA-seq. In contrast, for stimulation experiments, 5×10^5^ cells were seeded in a nucleon sphere 96-well round bottom plate in 200 μl RPMI1640 supplemented with 50 μg/mL gentamicin, 2 mM L-glutamine and 1 mM pyruvate. Then, *in vitro* stimulations were applied for either 3h or 24h using 1×10^6^ CFU/ml heat-killed *C. albicans blastoconidia* (strain ATCC MYA-3573, UC 820), 50 μg/ml heat-killed *M. Tuberculosis* (strain H37Ra, Invivogen) or 1×10^7^ heat-killed *P. Aeruginosa* (Invivogen) while incubating the cells at 37°C in a 5% CO_2_ incubator. After stimulations, cells were washed twice in medium supplemented with 0.04% BSA. Cells were then counted using a haemocytometer, and cell viability was assessed by Trypan Blue.

### Single-cell library preparation and sequencing

105 sample pools were prepared, each aimed to yield 1,400 cells/individual from 8 individuals (11,200 cells). In general, pools contained a mixture of both sexes and two different stimulation conditions. Each sample pool was loaded into a different lane of a 10x chip (Single Cell A Chip Kit for v2 or Single Cell B Chip Kit for v3 reagents). The 10x Chromium controller (10x Genomics), in combination with v2 (72 libraries) or v3 (33 libraries) reagents, was used to capture the single cells and generate sequencing libraries, according to the manufacturer’s instructions (document CG00052 and CG000183 for v2 and v3, respectively) and as previously described.^12^ Sequencing was performed with a 150 bp paired-end kit using a custom program (V2: 27-9-0-150, V3: 28-8-0-150) on the Illumina NovaSeq 6000 at BGI (Hong Kong).

### scRNA-seq alignment, preprocessing and QC

CellRanger v3.0.2 was used with default parameters to demultiplex, generate FASTQ files, align reads to the hg19 reference genome, filter both cell- and UMI barcodes and count gene expression per cell. To assign cells to one of the eight individuals in a lane, Demuxlet was used.^11^ The genotype information used by Demuxlet was previously generated as described in Tigchelaar et al.^70^ and was phased with Eagle v2.322 using the HRC reference panel and the Michigan Imputation Server. Only exonic variants with a MAF of at least 0.02 were used for demultiplexing. Subsequently, Souporcell v1.0^29^ was used to remove doublets coming from different individuals, by looking for the different genotypes within a single cell assignment. We limited the SNP calling to positions that were also used for demultiplexing.

Version 3.1 of the Seurat^71^ package was used for further quality control and processing. Due to mRNA capture differences between the v2 and v3 chemistries, a maximum mitochondrial gene content percentage of 7% and 15% was used, respectively. Cells with less than 200 detected genes were discarded, as were other low-quality cells (i.e. clusters of cells with a low number of expressed genes and a relatively high mitochondrial content, or missed, likely same-individual, doublets). Cells for each of the seven conditions and two chemistries were normalized separately by scaling the transcripts to 10,000 transcripts per cell, followed by log2 transformation. We then integrated the unstimulated data with each of the pathogen stimulations separately using Seurat’s IntegrateData function. We performed principal component analysis (PCA) and selected the first 20 PCs to identify the cell clusters using k-nearest neighbor clustering and visualized this in UMAP space (using the default settings). Cell types were assigned to each cluster based on marker gene expression, resulting in a set of six major cell types (*Fig. S1B*).

### Differential expression: mapping, pathway enrichment and module scoring

For each pathogen⍰timepoint combination, cell type and 10X chemistry, differential expression (DE) analysis was performed between the pathogen-stimulated and the untreated condition using the MAST implementation of Seurat.^30^ Testing was limited to genes with a log-fold change (LFC) >0.1 and with expression in at least 10% of the cells. We performed a meta-analysis for each cell type, taking the results of the v2 and v3 chemistries as input. Significance was determined by taking a Bonferroni-corrected p-value of <0.05 within the meta-analysis.

Per cell type, the resulting DE gene set was split in up- and downregulated genes after stimulation, which was then used as input for a pathway enrichment analysis with ToppFun, selecting the REACTOME database.^72^ To calculate statistical significance, the probability density function was used, selecting those pathways that had a BH-corrected p-value <0.05. To visualize the enriched pathways, we calculated the LFC in average gene expression in all pathogenCtimepoint conditions compared to the untreated condition and clustered these results using hierarchical clustering with the complete linkage method.

Calculation of pathway activity was done using the module score function of Seurat^73^, by calculating, per cell, the combined activity of a specific gene set annotated to be part of a pathway in the REACTOME database. This score was then averaged per donor for each condition and cell type.

### eQTL and re-QTLs: mapping and GWAS enrichment

The mapping of eQTLs was performed in a bulk-like and cell-type⍰specific manner. We limited the analysis to the top independent effects identified in the eQTLGen meta-analysis on 31,684 individuals, resulting in the testing of 16,987 possible SNPCgene pair combinations.^36^ These SNPCgene combinations identified by eQTLGen were the result of genome-wide *cis*-eQTL mapping of SNPs within a 100 kb distance to the gene midpoint, MAF >0.1, call rate >0.95 and Hardy-Weinberg equilibrium p-value >0.001. These 16,987 SNP⍰gene pairs were then further filtered to only include SNPs with a minor allele frequency (MAF) >0.1 or genes that were expressed in least three cells in our single-cell data. Filtering of SNP⍰gene combinations and mapping of eQTLs were done separately for each cell type and reagent version chemistry using the averaged, normalized gene expression values per individual, cell type and stimulation⍰timepoint combination. This was followed by a meta-analysis over the v2 and v3 chemistry outputs per cell type and stimulation⍰timepoint combination. eQTLs with a gene-level FDR < 0.05 were considered statistically significant, and a permutation-based strategy (n = 10) we had described before was used to control this FDR.^2^ Using the same parameters described above, but without eQTLGen SNP⍰gene pair filtering, we also performed a genome-wide *cis*-eQTL discovery analysis.

Next, we performed re-QTL mapping, confining ourselves to the total gene set of FDR < 0.05 significant eQTLs across all cell types and conditions. For this, we calculated the log-ratio of the averaged expression of the unstimulated condition and the stimulated condition per sample, cell type and chemistry, and then applied the same mapping strategy we used to identify regular eQTLs.

To determine whether eQTLs and re-QTLs were genetically inflated, eQTLgen lead eSNPs were matched to the top GWAS SNP per locus for each of the following immune-mediated disease GWAS studies: celiac disease^64^, type 1 diabetes^43^, multiple sclerosis^65^, inflammatory bowel disease^63^, candidemia susceptibility^67^ and rheumatoid arthritis^42^. For this, the LD between eSNPs and GWAS SNPs was calculated from genotypes of the 503 European individuals in the 1000g phase3 reference panel at R2 >0.8 using Plink 1.9-beta6.^74^ Lambda inflation was calculated using all GWAS p-values matched to the eQTL or re-QTL SNPs. To determine whether there is a difference in genomic inflation for those SNPs whose eQTL effect changes upon stimulation (re-QTLs), we compared the genomic inflation of the re-QTL SNPs with the non-re-QTL overlapping eQTL SNPs that were tested in both the unstimulated and relevant stimulation condition and significant in either. Using the different conditions and GWASes, specifically for the monocytes, the distributions of lambda values for the re-QTL and non re-QTL sets were compared using a two-sided Wilcoxon Rank Sum Test. This statistical testing was solely performed in monocytes, as this was the cell type with a strong pathogen response and the largest set of identified re-QTL SNPs, expecting largest effects on genomic inflation and allowing for the most robust genomic inflation analysis.

### Co-expression QTLs: mapping, pathway enrichment, TFBS and GWAS overlap

Co-expression QTL mapping was performed in the monocytes on a subset of SNPs and genes, selected based on their being: 1. DE and 2. a re-QTL in at least one of the stimulation-timepoint combinations; 3. expressed in at least 50% of the individuals (for each 10X chemistry tested). This selection resulted in 49 SNP⍰gene combinations for which we calculated the Spearman correlation with every other gene per individual and per stimulation⍰timepoint condition. A weighted linear model was used in which the genotype predicts the strength of the correlation between the two genes, using the square root of the number of cells as a weight. Analysis was performed separately for the different 10X chemistries, after which betas and standard errors were meta-analyzed. The statistical significance threshold was then determined using a permutation-based (n = 100) FDR approach. The most significant co-expression QTL p-value per stimulation⍰timepoint condition was then compared with the one coming from re-running the same permutations after randomly shuffling the genotype identifiers. This allowed us to calculate an eQTL gene-level FDR.^2^ An FDR < 0.05 was considered statistically significant. Separate thresholds were determined for each re-QTL SNPCgene combination and each stimulation-timepoint condition.

Pathway analysis was performed on the co-expression QTL genes associated with the selected eQTL gene per stimulation⍰timepoint combination using Toppfun with similar settings to those described in the ‘DE and pathway analysis’ section. Significant pathways (BH-corrected p-value < 0.05) were then ranked by p-value. The rankings of the pathways for each condition were then clustered using hierarchical clustering using the complete linkage method.

Transcription factor motif enrichment analysis was performed on the 3h stimulation outer join *CLEC12A* co-expressed gene set split by having either a more positive or more negative correlation with the minor versus major allele. For this, we took information from the TRANSFAC database Release 2020.2 v2^51^ and used g:Profiler (version e102_eg49_p15_7a9b4d6)^75^ with the g:SCS multiple testing correction method, applying a significance threshold of 0.05. Additionally, the *CLEC12A* co-expression QTL SNP rs12230244 and its accompanying (near-)perfect LD SNPs were overlapped with putative TFBSs, as defined by SNP2TFBS.^52^

Overlap of co-expression QTL SNPs (or SNPs within a 1 Mb window with LD >0.8) with disease-associated GWAS SNPs was determined by searching the GWAS catalog (https://www.ebi.ac.uk/gwas/) and an additional set of immune-related GWAS studies (celiac disease^64^, type 1 diabetes^43^, multiple sclerosis^65^, inflammatory bowel disease^63^, candidemia susceptibility^67^ and rheumatoid arthritis^42^).

### *CLEC12A* co-expression QTL validation and replication: SLE PRS interaction analysis, SLE scRNA-seq co-expression QTL analysis

Using the summary statistics of the SLE GWAS by Bentham et al.^62^, we calculated the PRS for SLE in 3,553 samples from the BIOS consortium using a custom Java program, GeneticRiskScoreCalculator-v0.1.0c, as described previously.^36^ Briefly, to account for LD between variants, our approach included a double clumping strategy where we first clumped variants within a 250 kb window and then within a 10 Mb window using an LD threshold R^2^ = 0.1. We then calculated the PRS for each individual by summing the products of the number of risk alleles and the GWAS effect-size (i.e. beta) for each SLE-associated variant. We constructed the PRS using a p-value threshold for the SLE GWAS of p < 5×10^−ss8^. The resulting PRS was scaled between 0 and 2 for compatibility with the eQTL mapping software. We then determined whether the co-expression between *CLEC12A* and an individual’s SLE PRS was modulated by SNP rs12230244. For this, we fitted a generalized linear model with and without the SLE PRS as an interaction term and determined how far the predicted model deviated from the true observation by taking the residuals of the observation. An F-test was then used to determine whether the distribution of the squared residuals with the SLE PRS as interaction term was significantly smaller than without, meaning that the SLE PRS interacts with the *CLEC12A* co-eQTL.

We used an independent cohort of SLE patients and healthy controls (GEO accession number: GSE174188) to replicate our findings of a clear enrichment for IFN-related genes within the co-expressed gene set of the *CLEC12A*-SNP rs12230244 co-expression-QTL. This cohort contained individuals of EUR and EAS descent, including healthy individuals (EAS: 18, EUR: 58) and individuals diagnosed with SLE (EAS: 58, EUR:59) who were not in an active disease state when samples were collected. For all individuals, PBMCs were collected and cryopreserved until further use. The SLE samples were collected through the California Lupus Epidemiological Study (CLUES) cohort. Healthy controls were collected at the UCSF Rheumatology Clinic and through the Immune Variation Consortium (ImmVar) in Boston. All UCSF samples were genotyped using the Affymetrix World LAT Array, while samples collected in Boston were genotyped using the Illumina OmniExpressExome Array. The Michigan Imputation Server was used for imputation with the Haplotype Reference Consortium version 1.1 reference. The samples collected at UCSF and Boston were processed using established protocols^11,27^. ScRNA-seq was performed using 10X Chromium Single Cell 3’ V2 chemistry, as described previously.^11^ Libraries were sequenced on the HiSeq4000 or NovaSeq6000 at a depth of 6,306-29,862 reads/cell. Freemuxlet was used to assign cells to individuals and, together with Scrublet^76^, for the identification of doublets. Marker gene expression was used to assign the major cell types. Only the monocytes were taken for this independent discovery analysis. Monocytes with less than 1500 UMIs were removed, as were donors with fewer than 200 cells remaining after applying this cutoff. Co-expression QTL analysis was performed as described in the co-expression QTL mapping paragraph above, but only testing the *CLEC12A*-SNP rs12230244 co-expression-QTL and doing this analysis separately in each cohort, ancestry and disease state (SLE versus healthy). A meta-analysis over the cohorts and ancestries was then performed, and pathway analysis using the REACTOME database was conducted to determine whether the IFN pathway was differently enriched in SLE compared to the healthy controls.

## Supporting information

All supplementary figures

## Data availability

Raw gene expression counts, eQTL and co-expression QTL summary statistics can be found under “Supplementary Data” at the website accompanying this paper (https://eqtlgen.org/sc/datasets/1m-scbloodnl.html). Processed (de-anonymized) scRNA-seq data, including a text file that links each cell barcode to its respective individual, has been deposited at the European Genome-Phenome Archive (EGA), which is hosted by the EBI and the CRG, under accession number EGAS… Gene expression and genotype data can be obtained and requested by filling in a short web form at https://eqtlgen.org/sc/datasets/1m-scbloodnl.html. This form is subsequently reviewed by a single Data Access Committee, who will be able to approve access to both the raw gene expression and genotype data within 5 working days (during the holiday season there might be a slight delay). Once the proposed research is approved, access to the relevant gene expression or genotyped data will be free of charge. Access to the genotype and gene expression data is facilitated via the HPC cluster of the UMCG and the EGA, respectively. Sample metadata (age, gender) is presented in **Table S1**.

## Code availability

The original code for Seurat v3.1^71^ (https://github.com/satijalab/seurat), Demuxlet^11^ (https://github.com/statgen/demuxlet), Souporcell v1.0^29^ (https://github.com/wheaton5/souporcell), Freemuxlet as part of the Popscle suite of statistical genetics tools (https://github.com/statgen/popscle), Scrublet^76^ (https://github.com/swolock/scrublet), and our in-house eQTL pipeline^2^ (https://github.com/molgenis/systemsgenetics/tree/master/eqtl-mapping-pipeline) can be found at GitHub. All custom-written code is made available via GitHub (https://github.com/molgenis/1M-cells).

## Acknowledgements

We are very grateful to all the volunteers who participated in this study and would like to thank K. McIntyre for proofreading the manuscript. L.F. and M.W. are supported by grants from the Dutch Research Council (ZonMW-VIDI 917.14.374 and ZonMW-VICI to L.F., NWO-VENI 192.029 to M.W.), and by an ERC Starting Grant, grant agreement 637640 (ImmRisk) and through a Senior Investigator Grant from the Oncode Institute. The Biobank-Based Integrative Omics Studies (BIOS) Consortium is funded by BBMRI-NL, a research infrastructure financed by the Dutch government (NWO 184.021.007). The images in Fig. 4f are created using Servier Medical Art, which we are thankful to for providing free online images.

## Author contribution

MW collected blood samples and generated the scRNA-seq data. RO, DV and HB performed bioinformatics and statistical analyses. RO, DV, HB and MW generated figures. HB built the website accompanying the manuscript. GG and JY performed and provided SLE scRNA-seq data for co-expression QTL replication analysis. MV and HW provided critical input for the statistical analyses. The BIOS consortium provided samples to conduct the SLE PRS co-expression QTL analysis. DV, MW and LF designed the study. RO, DV and MW wrote the manuscript and all other authors provided critical feedback. All authors discussed the results and commented on the manuscript.

## Ethics approval and consent to participant

The LifeLines DEEP study was approved by the ethics committee of the University Medical Centre Groningen, document number METC UMCG LLDEEP: M12.113965. All participants signed an informed consent from prior to study enrollment. All procedures performed in studies involving human participants were in accordance with the ethical standards of the institutional and/or national research committee and with the 1964 Helsinki declaration and its later amendments or comparable ethical standards.

