## Supplementary figures and images for "Single-cell RNA-sequencing reveals widespread personalized, context-specific gene expression regulation in immune cells"

### Figure S1

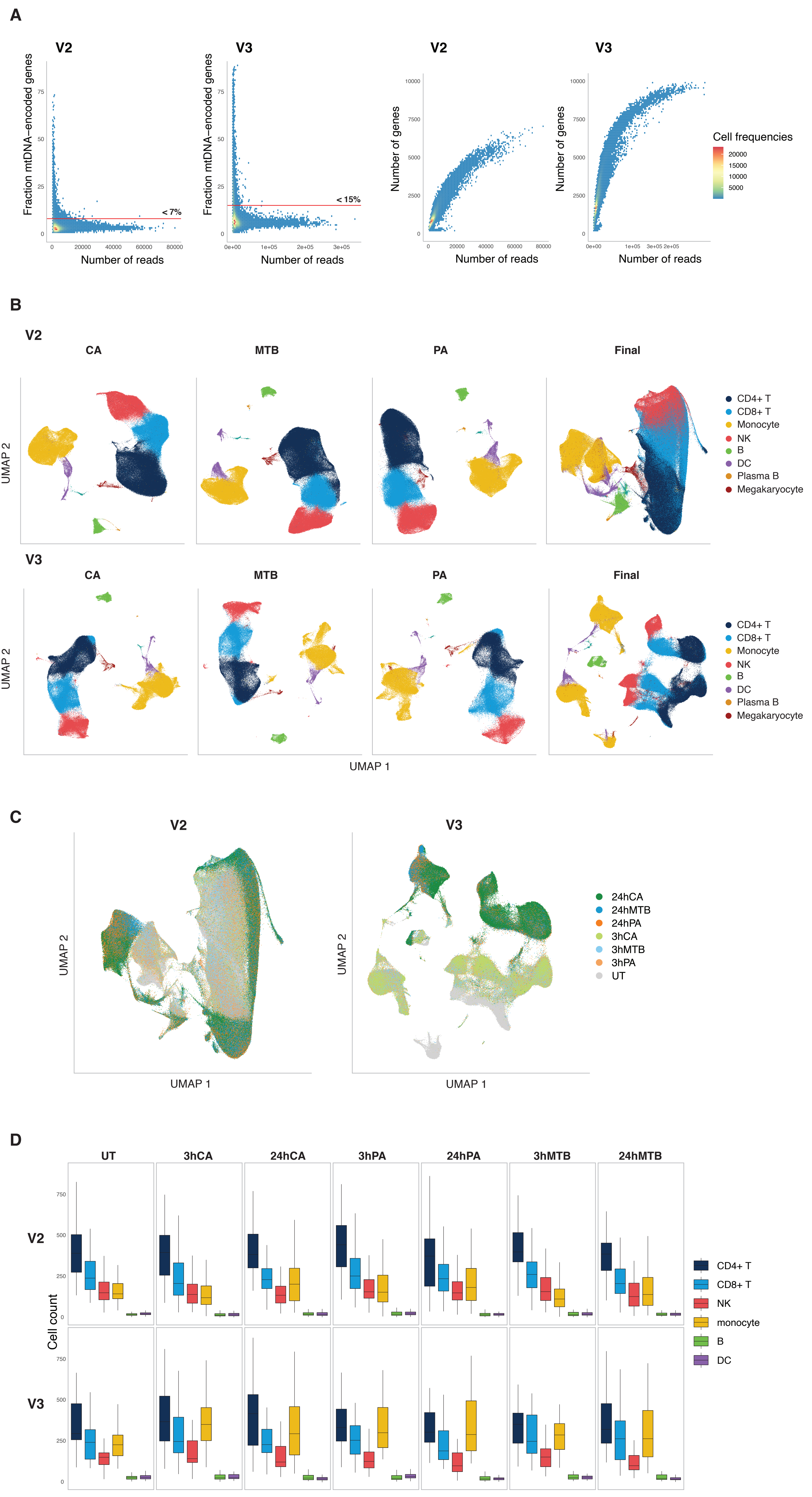

### Figure S2

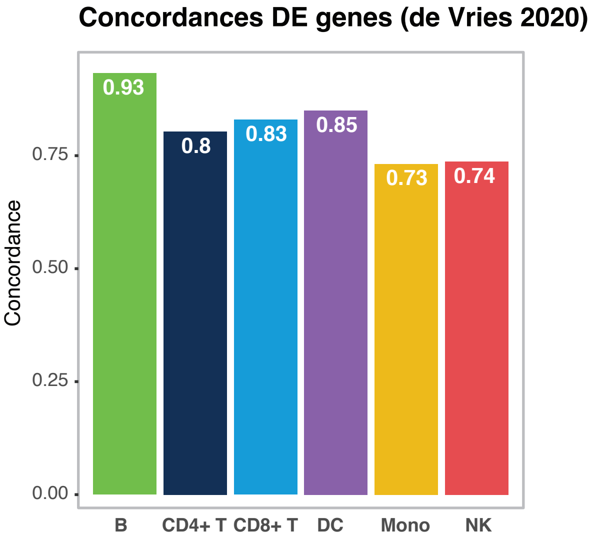

### Figure S3

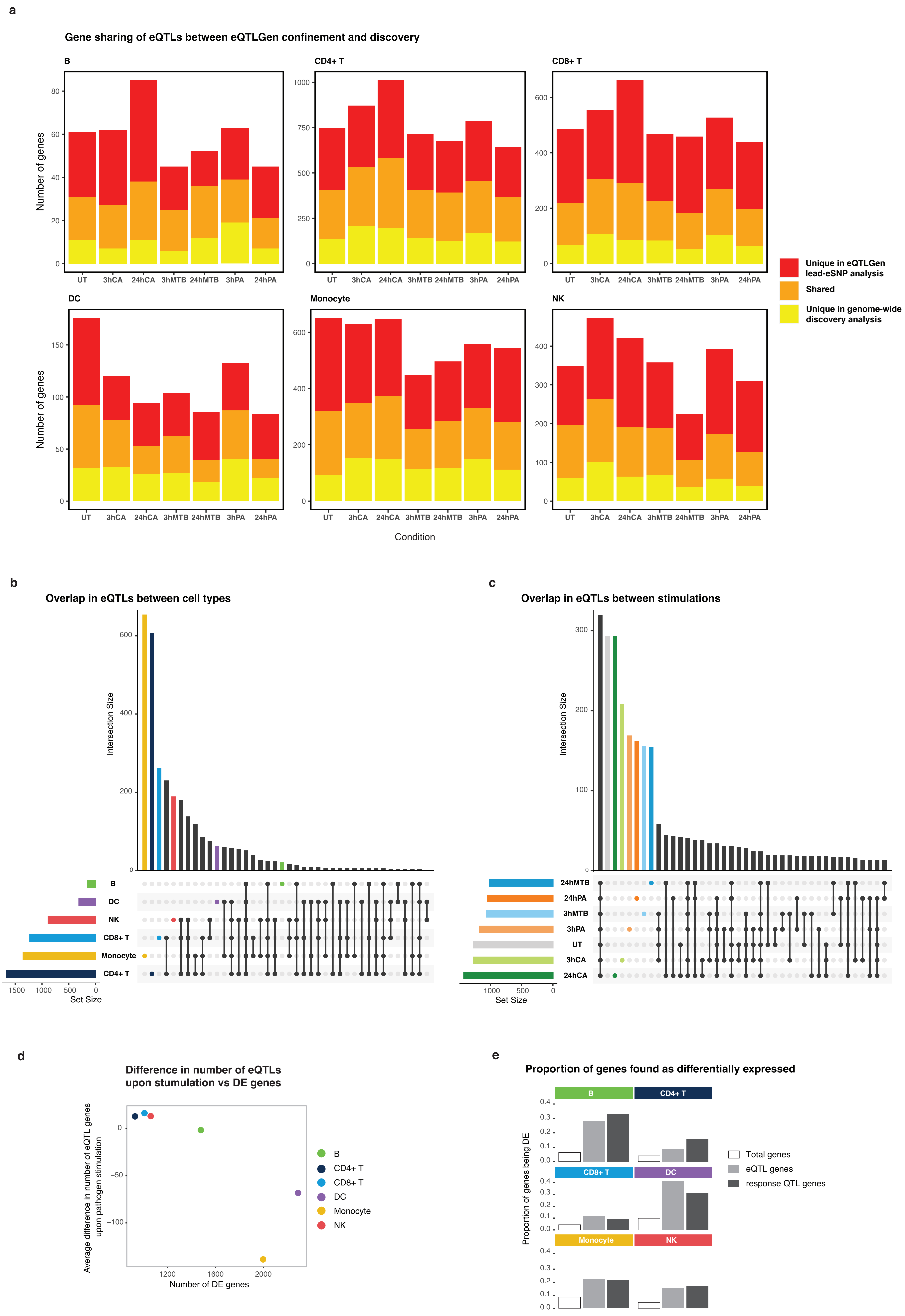

### Figure S4

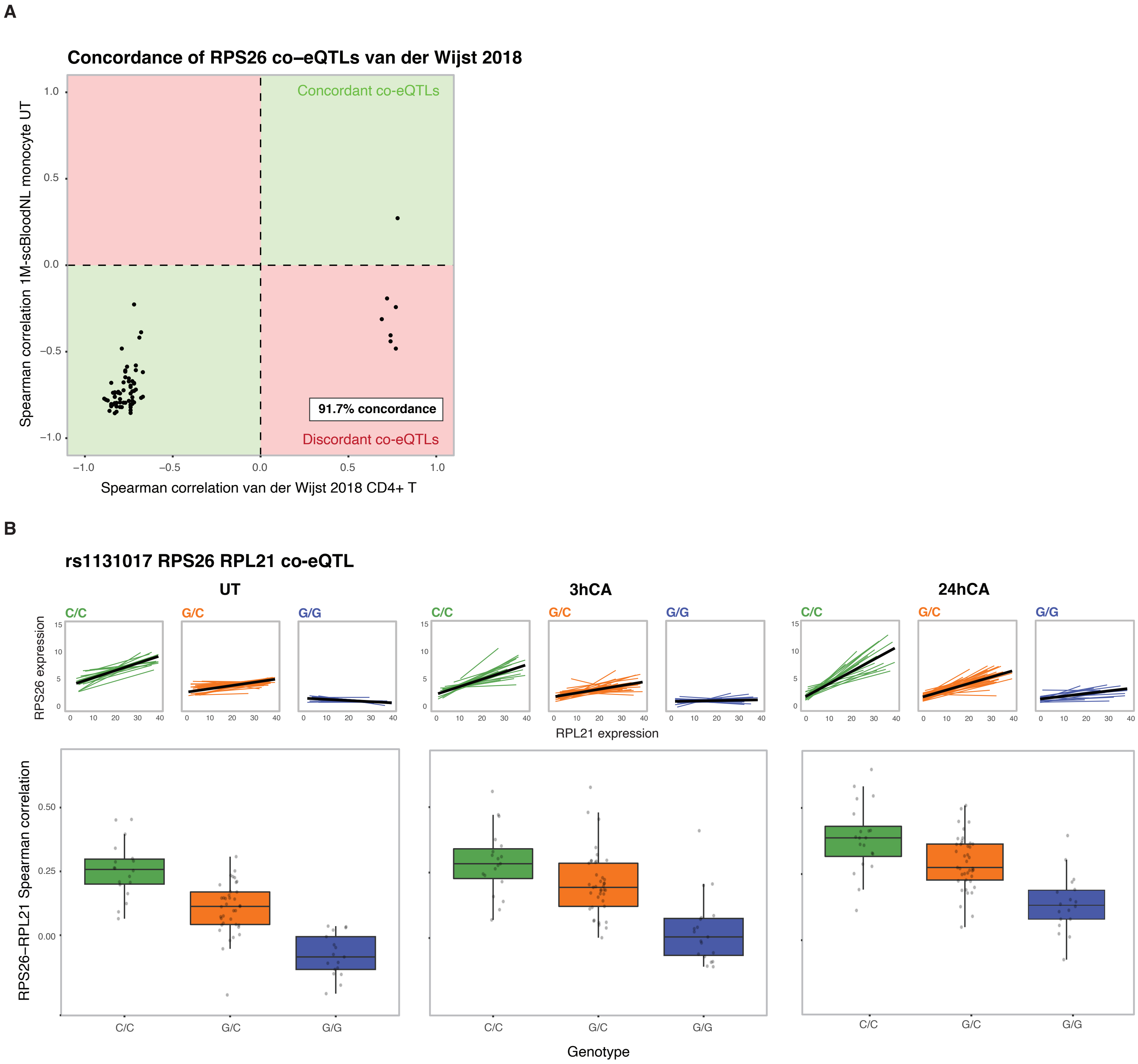

### Figure S5

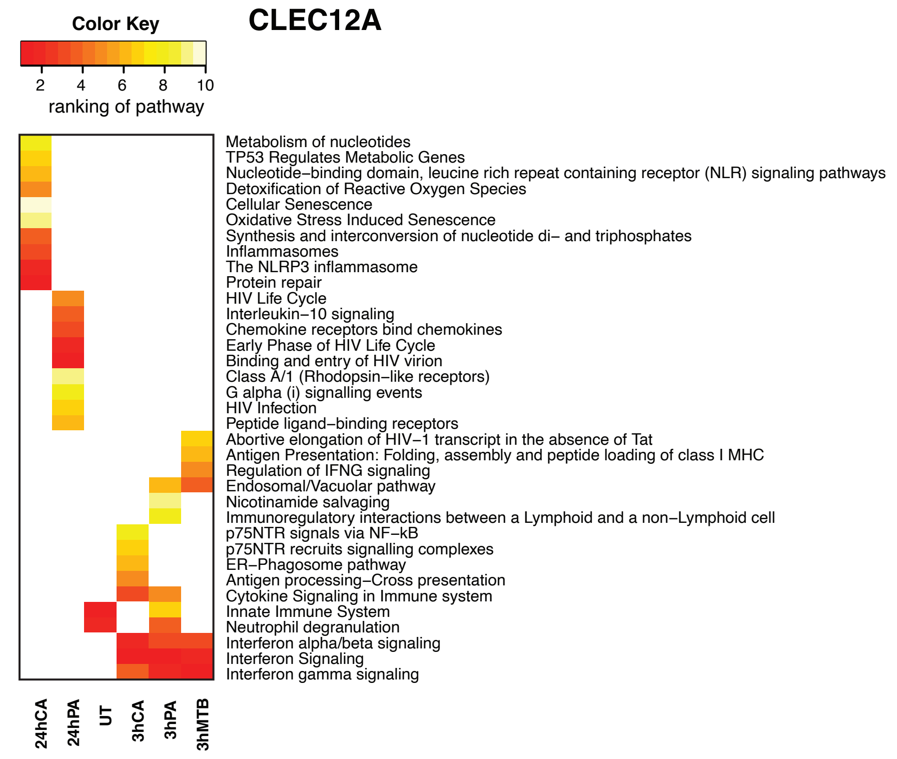

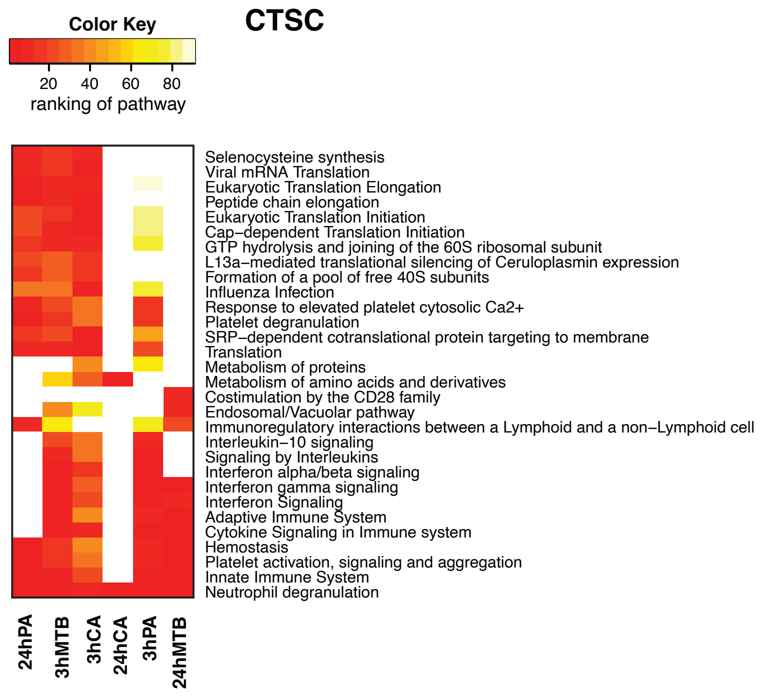

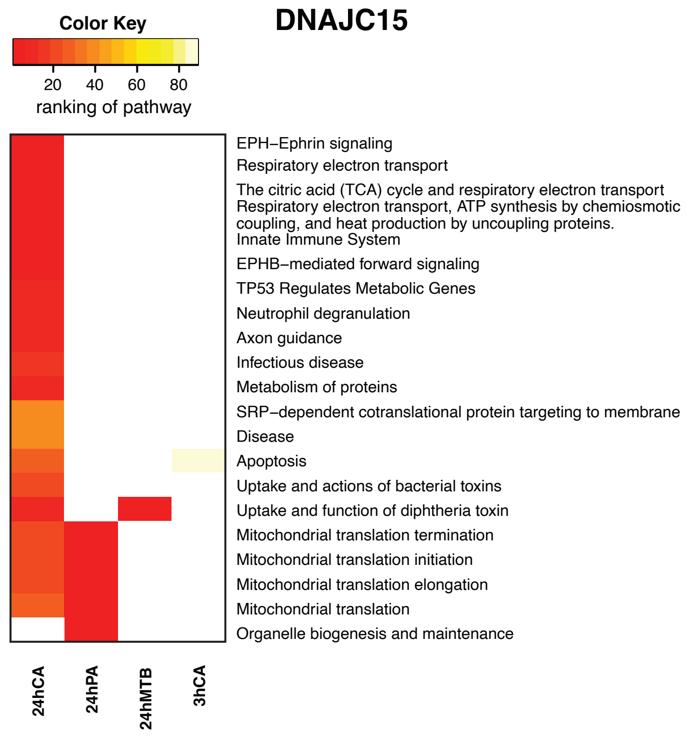

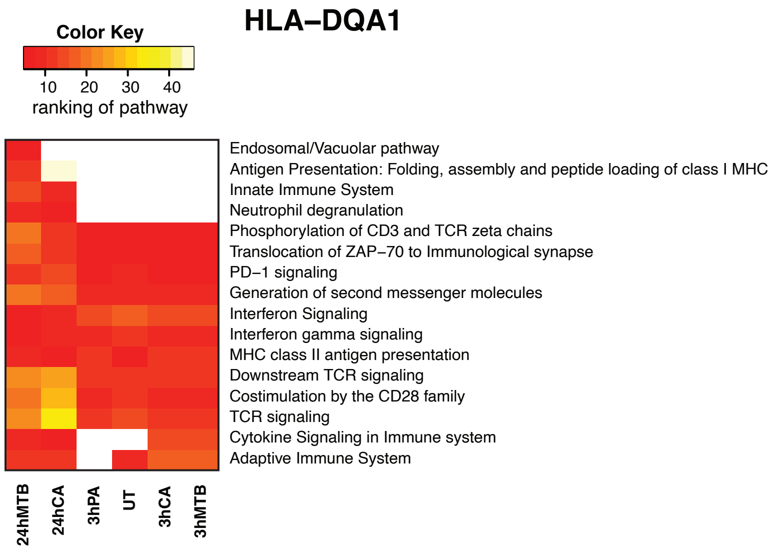

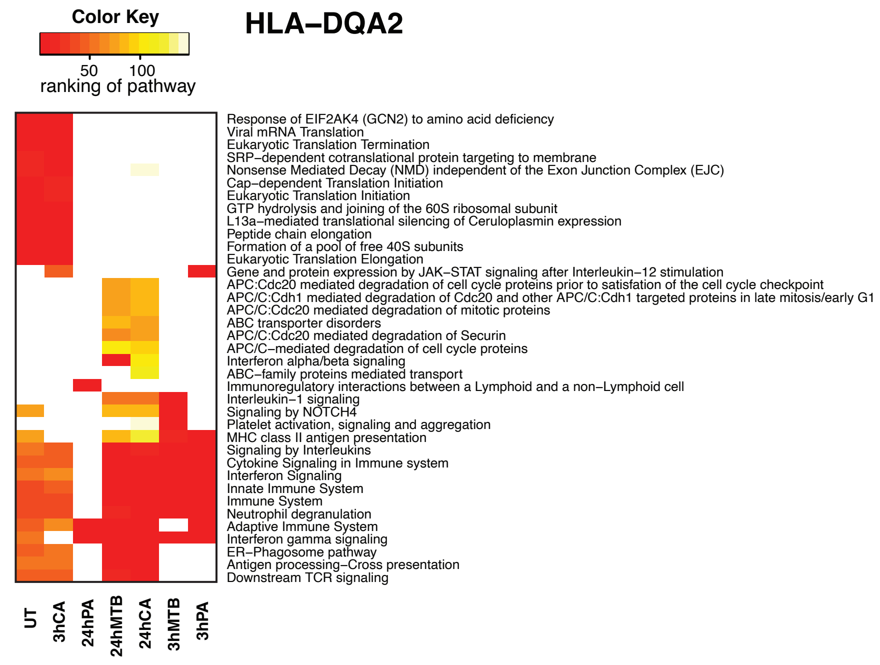

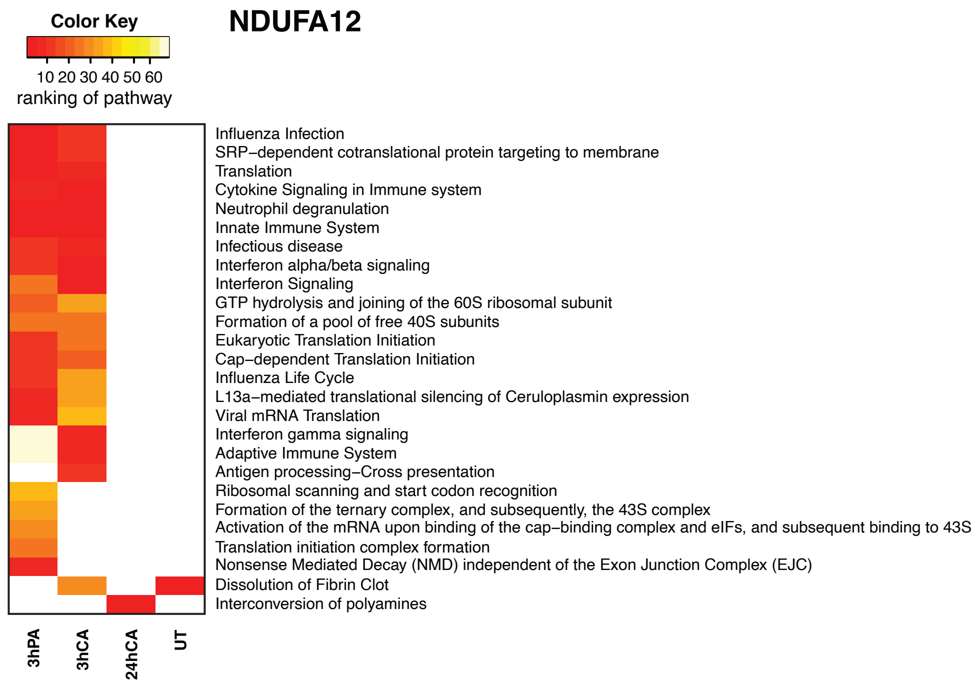

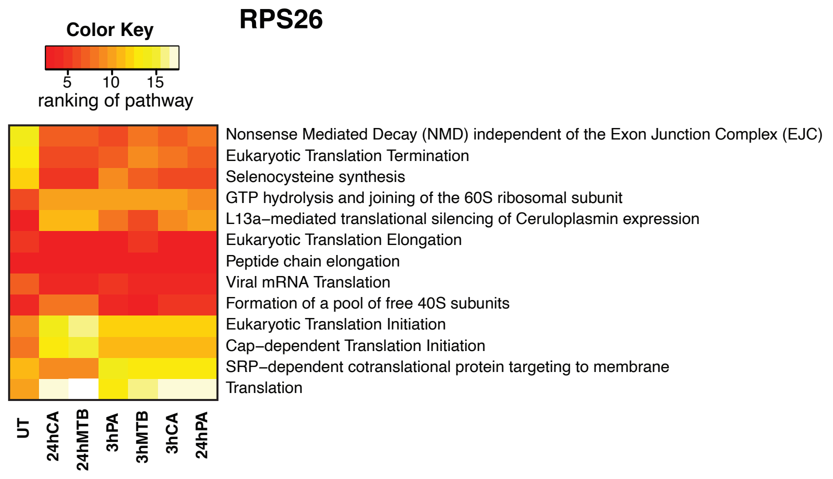

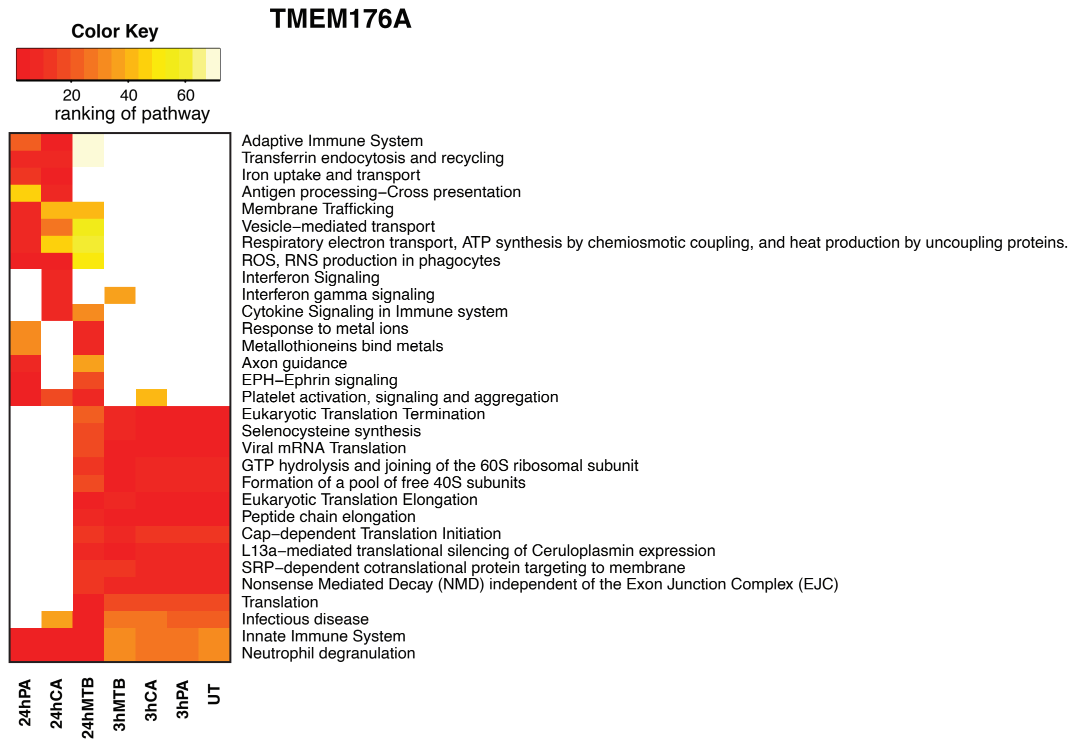

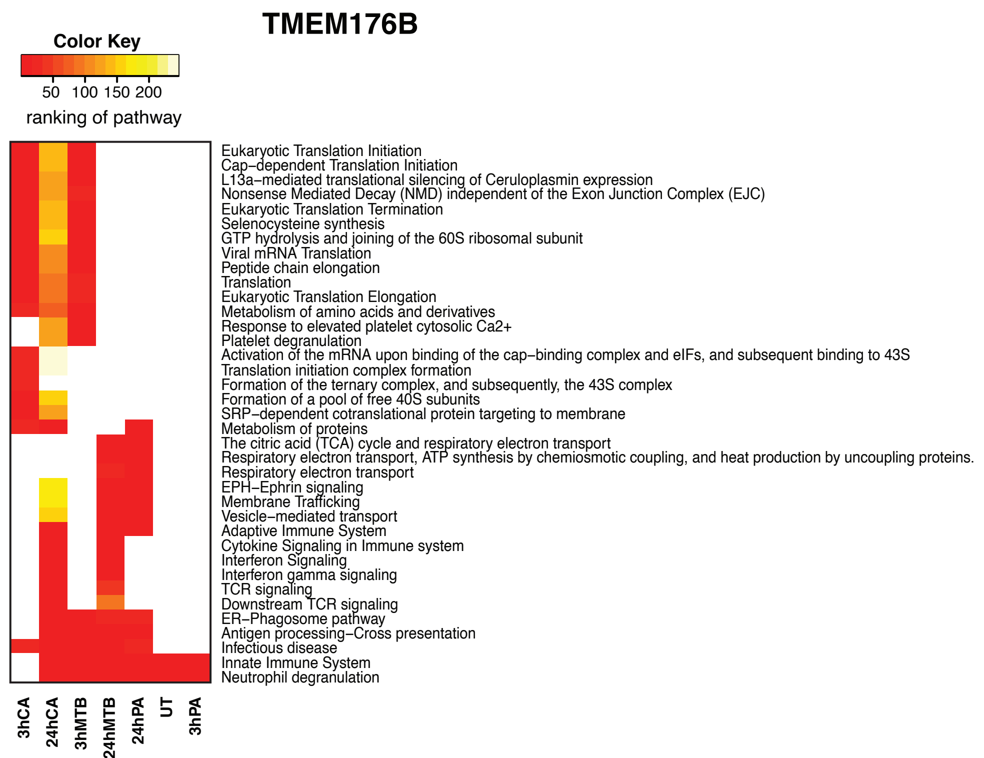
