## Supplementary material for "Single-cell RNA-sequencing reveals widespread personalized, context-specific gene expression regulation in immune cells": All supplementary figures

**Supplementary Information**

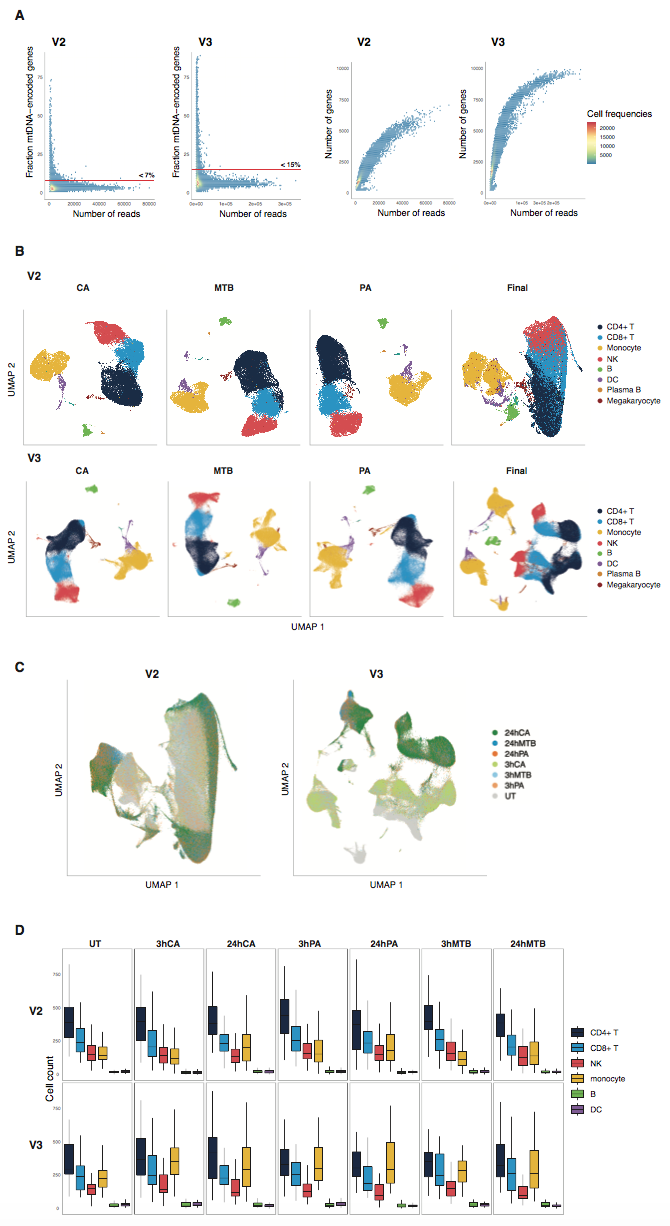

***Figure S1. Dataset characteristics***

***a.*** *The proportion of mitochondrial genes (left two panels) or number of expressed genes (right two panels) per cell (y-axis) against the number of reads per cell (x-axis), split per sequencing chemistry (v2 or v3). Cell density is indicated by color in each graph, going from blue to red for low to high cell numbers, respectively. The red line in the left two panels indicates the QC threshold used for removing cells with a high mitochondrial gene fraction.* ***b.*** *The UMAP plots per stimulation and chemistry. Each dot represents a single-cell and the color indicates the assigned cell type.* ***c.*** *The integrated UMAP per chemistry where all cells are combined.* ***d.*** *Boxplots (showing median, 25th and 75th percentile, and 1.5 x the interquartile range) representing the cell type proportions per individual, split by stimulation-timepoint combination and chemistry. Colors represent the cell types.*

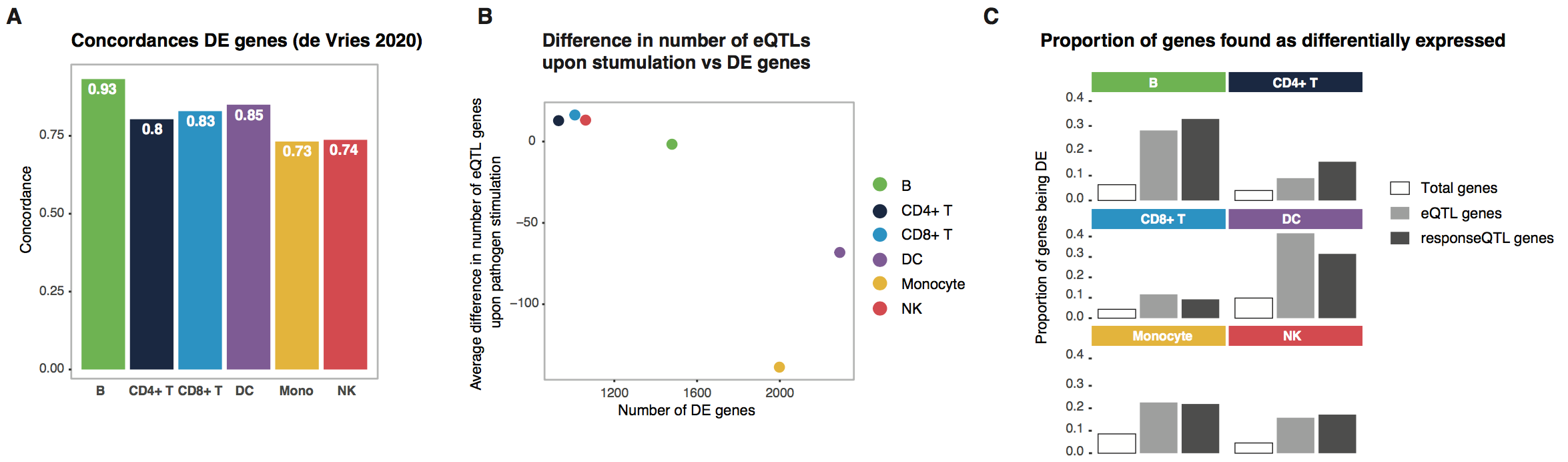

***Figure S2. Concordance differential expression results with literature***

***a.*** *Bar plot showing the concordance of the identified differentially expressed (DE) genes in this study with those identified in de Vries et al. 2020. Each bar represents a different cell type.*

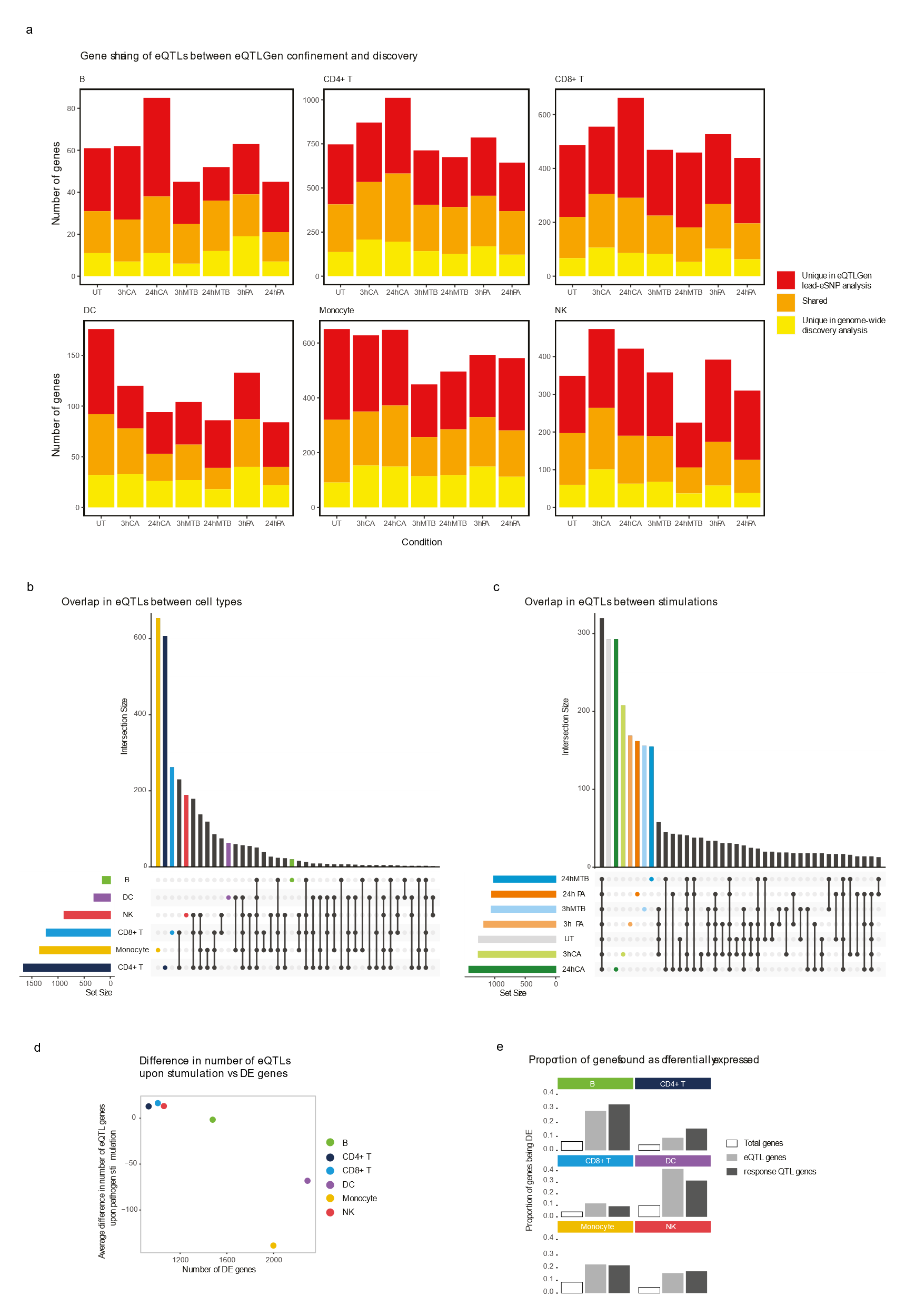

***Figure S3. eQTL characteristics***

***a.*** *Stacked bar plots showing the total number of unique eQTLs per stimulation-timepoint combination within the eQTLGen lead-eSNP confined eQTL analysis and the genome-wide cis-eQTL discovery analysis. The red color shows eQTLs that were uniquely identified in the eQTLGen lead-eSNP confined analysis, orange is shared across both analyses and yellow shows eQTLs that are unique to the genome-wide cis-eQTL discovery analysis.* ***b.*** *Overlap of eQTL effects between cell types and* ***c.*** *stimulation-timepoint combinations. Each bar represents the number of eQTLs found for that group, which is indicated by the dots underneith the bar. Coloured bars show eQTLs that are unique to one group and black bars are a combination of different groups.* ***d.*** *Dot plot showing the difference in number of identified eQTLs upon stimulation (y-axis) compared to the number of DE genes (x-axis). Each dot represents a different cell type, shown by the color.* ***e.*** *Bar plots showing the proportion of genes that were identified as DE per cell type. The three bars represent the complete set of tested genes (white), the complete set of genes with at least one eQTL (gray) and the complete set of genes with at least one response QTL (dark gray).*

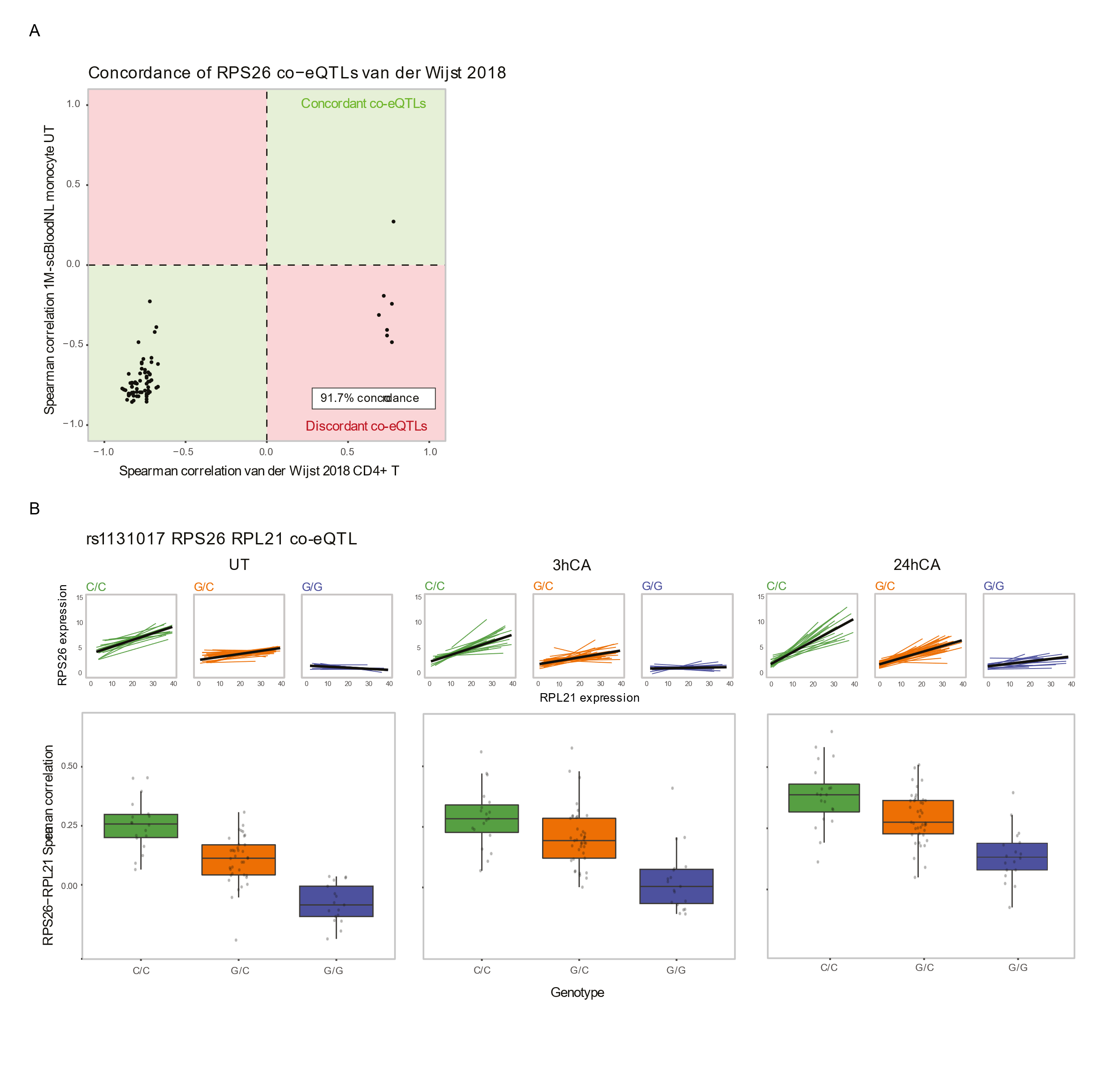

***Figure S4. RPS26 co-expression QTLs***

***a.*** *Concordance plot comparing Spearman correlations of RPS26 co-expression QTLs in CD4+ T cells from van der Wijst et al. 2018 (x-axis) compared to RPS26 co-expression QTLs in monocytes in the untreated condition of this study. Each dot represents a different co-expression QTL. Dots in green quadrants are concordant and dots in red quadrants are discordant.* ***b.*** *The co-expression QTL of RPS26 and RPL21, mediated by rs1131018, across different stimulation-timepoint combinations. The top graphs show the individual co-expression, with each colored line representing one individual and the black line showing the regression line across all points. Boxplots (showing median, 25th and 75th percentile, and 1.5 x the interquartile range) representing the Spearman correlations per individual, split by genotype group. Each dot represents the Spearman correlation between RPS26 and RPL21 for one individual within that genotype group. Colors represent the three genotype groups for rs1131018.*

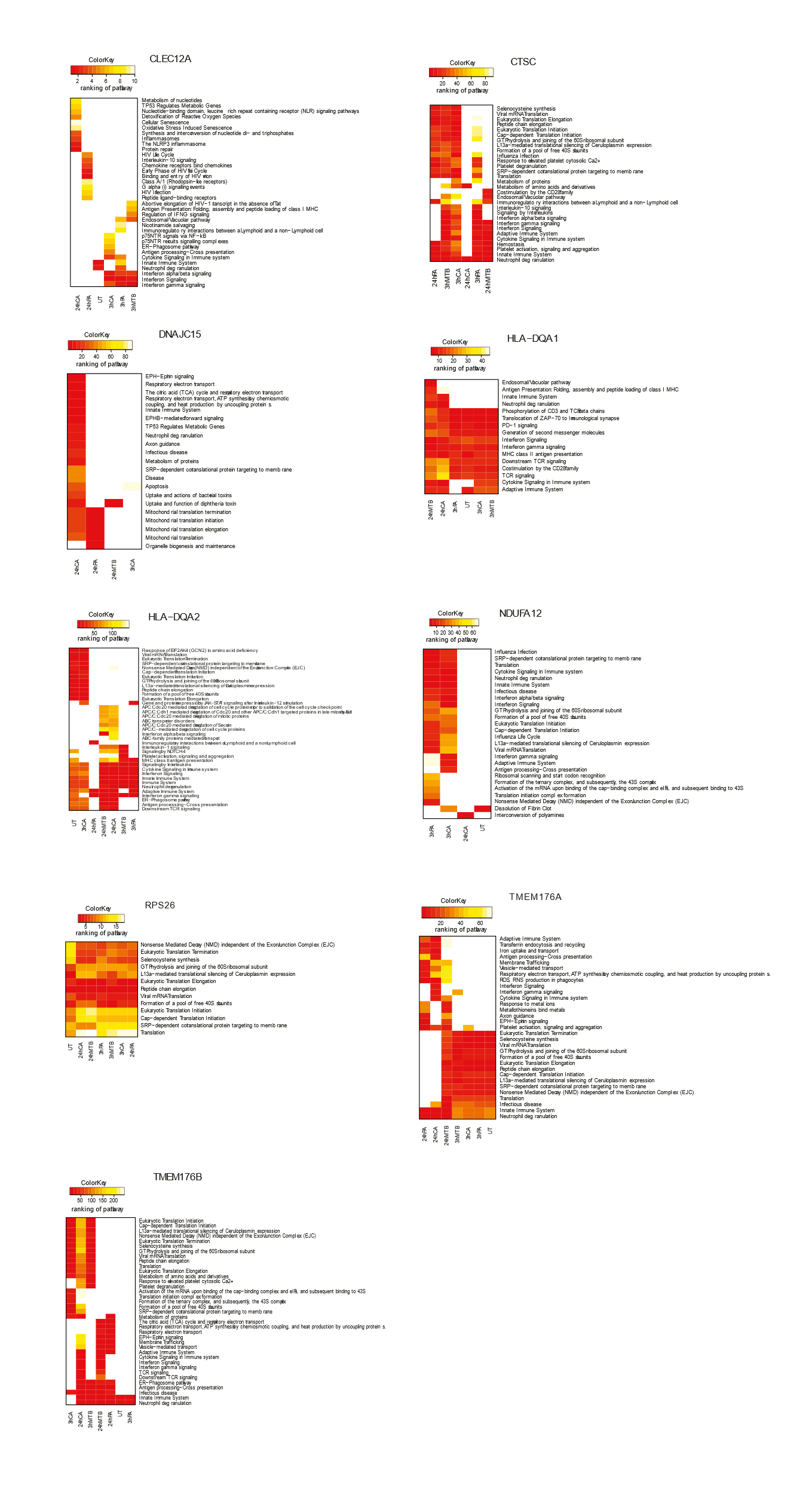

***Figure S5. Pathway enrichment analysis of co-expression QTL gene sets***

*Heatmaps showing the enrichment ranks of the pathways associated with the set of co-expressed genes affected by each co-expression QTL gene. Darker colors represent lower ranks, i.e. stronger enrichment, and lighter colors represent higher ranks, i.e. less enrichment.*

***Table S1. Sample metadata***

***Table S2. DE analysis***

***Table S3. Pathway analysis***

***Table S4. eQTLgen lead-eSNP eQTL analysis***

***Table S5. Genome-wide eQTL analysis***

***Table S6. Response-QTL analysis***

***Table S7. Genomic inflation analysis***

***Table S8. Co-expression QTL analysis in monocytes***

***Table S9. Co-expression QTL analysis SLE cohort in monocytes***
